# Targeted hypermutation of putative antigen sensors in multicellular bacteria

**DOI:** 10.1101/2023.09.22.558884

**Authors:** H. Doré, A. R. Eisenberg, E. N. Junkins, G. E. Leventhal, Anakha Ganesh, O. X. Cordero, B. G. Paul, D. Valentine, M. A. O’Malley, E. G. Wilbanks

## Abstract

Diversity-generating retroelements (DGRs) are used by bacteria, archaea and viruses as a targeted mutagenesis tool. Through error-prone reverse transcription, DGRs introduce random mutations at specific genomic loci, enabling rapid evolution of these targeted genes. However, the function and benefits of DGR-diversified proteins in cellular hosts remains elusive. We find that 85% of DGRs from one of the major monophyletic lineages of DGR reverse transcriptases are encoded by multicellular bacteria, which often have two or more DGR loci in their genomes. Using the multicellular purple sulfur bacterium *Thiohalocapsa* sp. PB-PSB1 as an example, we characterized nine distinct DGR loci that cumulatively lead to more than 10^294^ possible different combinations of target polypeptides. With environmental metagenomes from individual *Thiohalocapsa* aggregates, we show that most of PB-PSB1’s DGR targets are diversified across its biogeographic range, with spatial heterogeneity in the diversity of each locus. In PSB1 and other members of this lineage of cellular DGRs, diversified target genes are associated with NACHT-domain anti-phage defenses and putative ternary conflict systems previously shown to be enriched in multicellular bacteria. We propose that these DGR-diversified target genes act as antigen sensors that confer a form of adaptive immunity to their multicellular consortia. These findings have implications for the emergence of multicellularity, as the NACHT-domain anti-phage systems and ternary systems share both domain homology and conceptual similarities with the innate immune and programmed cell death pathways of plants and metazoans.

**Significance:** To defend themselves against predators, bacteria employ a wide range of conflict systems, some of which are enriched in multicellular bacteria. Here, we show that numerous multicellular bacteria use related diversity-generating retroelements (DGRs) to diversify such conflict systems. Error-prone reverse transcription in DGRs introduces random, targeted mutations and rapid diversification. We used *Thiohalocapsa* PB-PSB1, a member of multicellular bacterial consortia, as a model to study this association between conflict systems and DGRs. We characterized the natural diversity of PB-PSB1 DGRs and propose they function as hypervariable antigen sensors. The accumulation of such DGR-diversified defense systems in multicellular bacteria emphasizes the fitness advantage of a rapidly diversifying immune system for the evolution of multicellularity.

## Introduction

In the evolution of life, a handful of major transitions mark turning points in the emergence of complexity (1, 2). Such evolutionary transitions, including from genes to genomes and from single cells to multicellular organisms, represent a shift in the nature of the individual and require cooperation amongst previously distinct entities. Explaining the emergence of cooperation remains a major challenge in understanding these transitions. Why should a cell sacrifice its individual interests in favor of the collective? Kin selection and inclusive fitness are commonly invoked to explain the evolution of cooperation: both theory and empirical evidence indicate that the clonality of the group is critical to minimizing conflict and paving the way for multicellularity (3, 4). However, close physical association in a group with few genetic differences also creates conditions for an infectious epidemic. In formulating his social evolutionary theory, Hamilton recognized that, given this lack of genetic diversity, disease could represent a major constraint on the emergence of multicellularity (5). How then do nascent multicellular forms balance the risks of infection with the benefits of cooperation?

While innate immunity was classically thought to have emerged amongst multicellular metazoans, discoveries many, novel defense systems have revealed the bacterial origins of numerous key components of innate immunity (6–11). Intriguingly, several of these recently discovered putative defense systems were found to be particularly enriched in bacteria with a multicellular lifestyle (8, 12–14). The latter systems were hypothesized to mediate immune-like recognition of invaders and trigger programed cell death to protect their clonal kin. Protein and carbohydrate binding domains were proposed to act as sensors of an invading phage while effector domains commonly included trypsin- or caspase-type peptidases. In some of these systems, short non-enzymatic adapter domains fused to sensor and effector domains (“effector associated domains” or EADs) are thought to mediate the assembly of a protein complex akin to the eukaryotic apoptosome and inflammasome, and in some cases are homologous to eukaryotic domains with this function (e.g. death-like and TRADD-N domains) (13). The formation of these protein complexes, and the resulting programmed cell death, is thought to be tightly regulated by the activity of associated NTPase or protein kinase domains (8, 15).

In several studies, these novel systems were found to be associated with diversity-generating retroelements (DGRs) (8, 13) or with the ligand binding domains DGRs commonly diversify (14). Discovered twenty years ago, DGRs are a unique class of retroelements found in bacteria, archaea, and viruses that generate massive sequence variation at specific protein-coding regions by mutagenic retrohoming (16–20). After transcription of a template repeat (TR) into a TR-RNA, the error prone reverse transcriptase (RT) reads the TR-RNA and incorporates random nucleotides at adenine sites to produce a mutated TR-cDNA (21). The TR-cDNA then recombines with the host genome, replacing a matching variable repeat (VR) region in the targeted gene (Fig. 1A). This process thus creates variability in the target gene at positions of the VR corresponding to TR adenines. The TR DNA sequence remains unaltered, enabling repeated rounds of diversification. The VR is most often located within a C-type lectin (CLec) fold (22–24), a ligand binding domain that can accommodate high amino-acid diversity while maintaining protein stability (22, 25).

**Figure 1.**
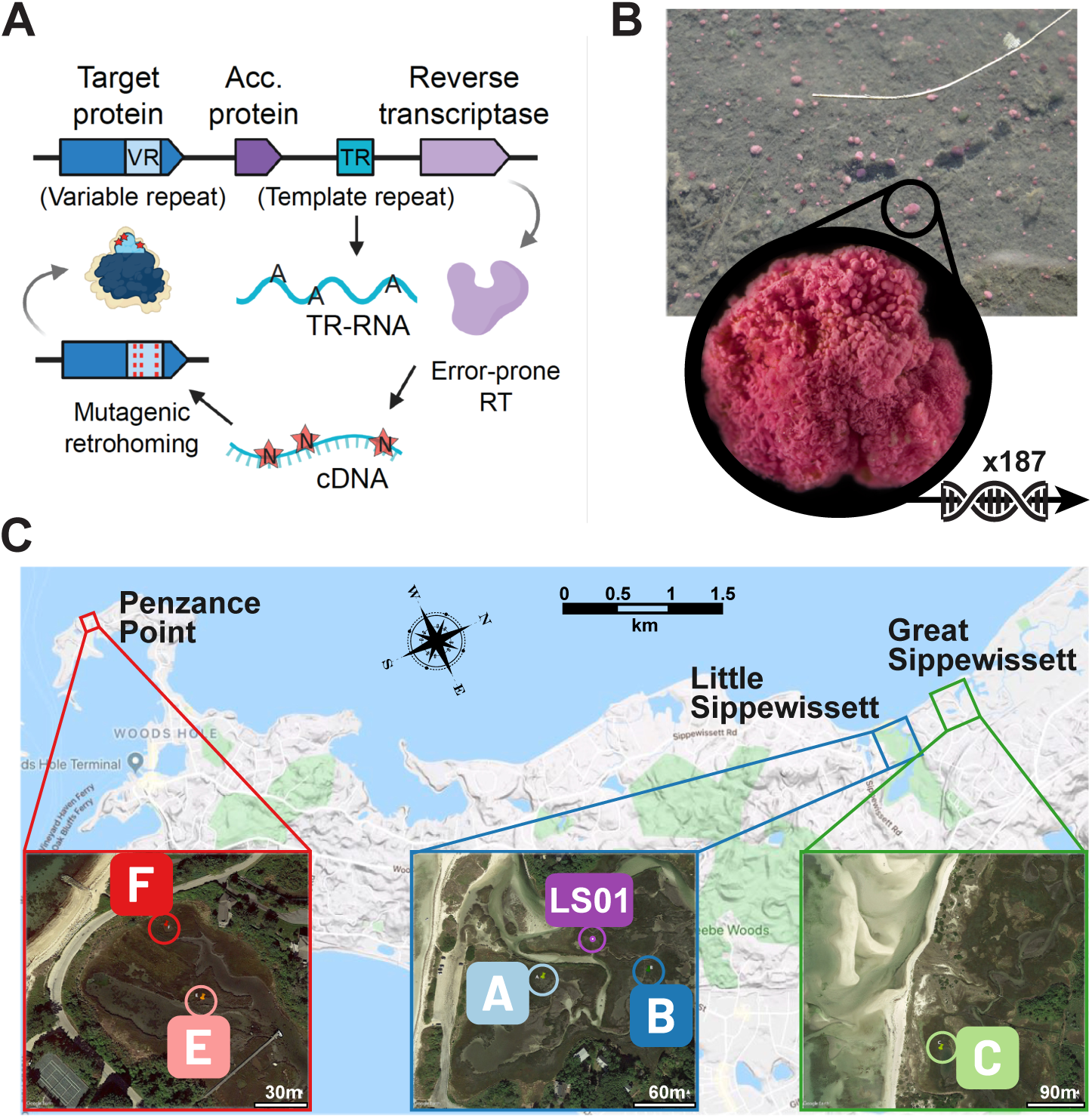
Schematic of DGR mechanism and sampling locations for the “pink berry” bacterial consortia. **(A)** Mutagenic retrohoming introduces mutations within the variable repeat (VR, light blue) of a target protein (dark blue). The error-prone reverse transcriptase (RT, light purple), which complexes with accessory protein(s) (acc, dark purple), introduces random nucleotides at adenine positions in the template repeat (TR, teal) generating a hypervariable TR cDNA that recombines into the VR of the target. **(B)** Individual pink berry aggregates were sampled in three salt marshes (Little Sippewissett, LS; Great Sippewissett, GS and Penzance Point, PP) and sequenced with both long read (n=3, site LS01) and short read (n=184, sites A, B, C, E and F) technologies at 6 sites across their geographic range (**C)**.

While much of the DGR mechanism has been elucidated, the function and benefit of DGRs for bacteria and archaea remain mysterious. Since the discovery of the first DGR as a mechanism for tropism switching in *Bordetella* bacteriophage (16, 17), very few DGR targets have been functionally characterized. This is particularly true for cellular (i.e. non-viral) DGRs: DGR diversification has been molecularly confirmed only in *Legionella pneumophila* (26), and the molecular structure was determined for a single cellular DGR target protein from *Treponema denticola* (23). In both these cases, the target genes are outer membrane lipoproteins thought to diversify the cell surface of these known pathogens. All other DGR cellular targets have been only computationally predicted. By similarity to proteins or protein domains in databases, several categories of functions have been proposed for these targets, including a role in interaction with the host in parasites or epibionts amongst CPR and DPANN organisms (19) and in gut and oral bacteria (27, 28), a role in regulation or signal transduction (19, 29, 30), or a role in defense against viruses (20, 28, 31, 32). In multicellular cyanobacteria, Vallota-Eastman *et al.* argued that DGR targets were likely involved in signal transduction, with unknown effects that could potentially range from regulation of cellular homeostasis to cellular differentiation or programmed cell death (30). Hence, the function of DGR target proteins, particularly for non-pathogenic microbes, remains elusive (20, 24).

The association of novel defense systems with DGRs in multicellular bacteria suggests an intriguing possibility: the diversification of antigen sensors, analogous to the somatic hypermutation of the vertebrate adaptive immune system (33). Here, we explore this possibility using multicellular bacterial consortia, the “pink berries” from salt marshes near Woods Hole, MA (USA), which were reported to have this DGR-conflict systems association in two previous studies (8, 13). These millimeter-sized aggregates, found at the water-sediment interface (Fig. 1B), offer a model to study bacterial multicellularity across time and space (34). They have a relatively simple species composition, with a few species accounting for most of the cells (35). Among these dominant species are a purple sulfur bacterium (*Thiohalocapsa* PB-PSB1) and a sulfate-reducing bacterium (PB-SRB1), which interact in a cryptic sulfur cycle (35). Although these bacteria are uncultivated, the genome of PB-PSB1 has recently been assembled into a single circular contig from long-read metagenomes, revealing its large size (8 Mb), and enrichment with transposable elements (36). This completed genome allows for a detailed exploration of DGR systems in a likely-obligatory member of multispecies cellular consortia.

We find that the multicellular bacterium *Thiohalocapsa* PB-PSB1 hypermutates putative antigen sensors using a suite of distinct DGR loci. *Thiohalocapsa* PB-PSB1 genome encodes the unusually high number of nine distinctly organized DGR loci that are diversified across PB-PSB1’s geographic range. These DGRs belong to one of two major monophyletic lineages of DGR reverse transcriptases from bacteria and archaea (clade 5 *sensu* (20)), and our in-depth analysis indicates that over 82% of the DGRs in this clade are encoded by multicellular bacteria. We show how mobile genetic elements are involved in the dynamics of DGR loci in PB-PSB1, and use metagenomics data from 187 independent aggregates sampled at six sites across the pink berries’ geographic range to elucidate whether all nine DGR loci are diversifying in natural conditions (Fig. 1B and 1C). Our careful analysis of the genomic context of clade 5 DGRs leads us to propose that they diversify the sensors of bacterial immune systems, and we discuss implications of this function in light of the selective pressures acting on multicellular life forms.

## Results

### Detection of nine DGR in *Thiohalocapsa* PB-PSB1

Nine DGR loci were found throughout the genome of *Thiohalocapsa* PB-PSB1, with a total of 15 identified target genes (Fig. 2, SI Dataset S1). These nine loci were grouped into four distinct classes based on both the phylogenetic relationships of the DGR reverse transcriptase (RT) genes and the alignment of template and variable repeat pairs (TR-VR) (Fig. 2B; SI Appendix Fig. S1). Each class includes a single locus with the complete repertoire of functional components (a reverse transcriptase, a template repeat, and an accessory protein), in addition to ‘degenerate’ loci where key machinery is either missing or pseudogenized (Fig. 2).

**Figure 2.**
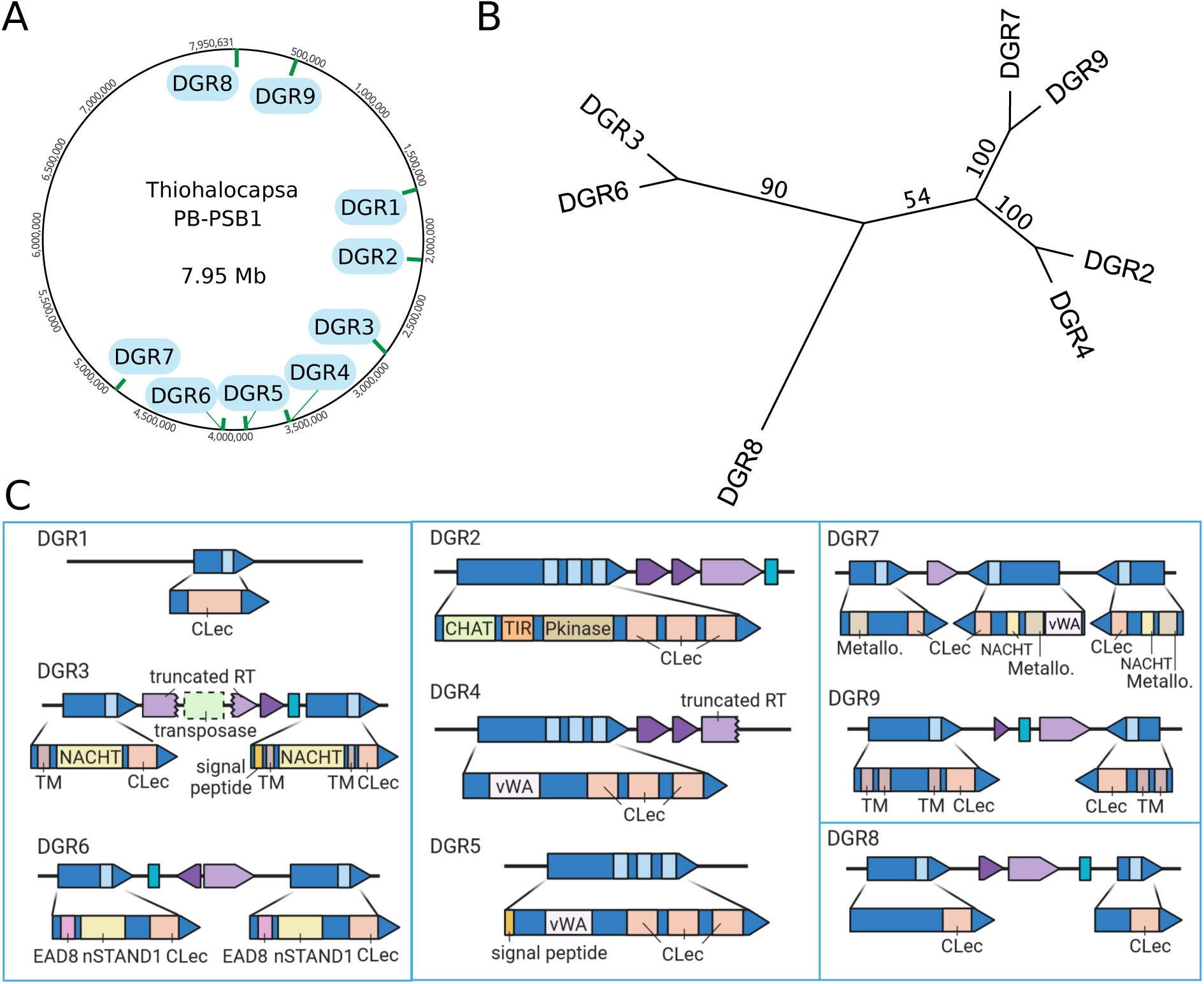
*Thiohalocapsa* PB-PSB1 has nine DGR loci spread across its 7.95 Mb circular, metagenome-assembled genome **(A)**. Loci were categorized according to the maximum likelihood phylogeny relationships of their reverse transcriptase genes **(B)** and matching template-variable repeats. The four distinct classes are indicated by blue boxes in **(C)** where each DGR locus is shown with annotated domains for target proteins. Colors scheme corresponds to Fig. 1A. Domain abbreviations include: C-terminal lectin domain (CLec); transmembrane domain (TM); von Willebrand domain (vWA); NACHT domain (PF05729); nSTAND1 domain (nSTAND1); CHAT domain (PF12770); Toll/interleukin-1 receptor domain (TIR, PF13676); protein kinase domain (Pkinase; PF00069); 3’,5’-cyclic AMP phosphodiesterase (Metallo; PF00149).

Together, the 15 DGR target proteins comprise 21 VRs with 521 potential variable positions that affect 325 different codons. The variable positions were almost exclusively found in the codon’s first or second position (often both), a pattern that maximizes the number of possible protein sequences (SI Appendix Fig. S1, SI Dataset S2). As was noted in other organisms, the composition of targeted codons also prevents the adenine-directed variation from creating nonsense codons (SI Dataset S2) (22). In PB-PSB1, DGR mutagenesis can yield from 10^9^ to 10^16^ different polypeptides per VR, and up to 2 × 10^^45^ different polypeptides per target (SI Dataset S2). This gives an astronomical total of 1.5 × 10^294^ possible sequence combinations at the protein level when accounting for all targets in the genome.

### Multicellular bacteria encode multiple phylogenetically related DGR reverse transcriptases

The seven DGR RTs from PB-PSB1 all belong to clade 5 (*sensu* (20)), one of two major monophyletic groups of DGR RTs originating from bacteria and archaea (as opposed to phage). Exploring the phylogeny of clade 5 RTs, we found that 82% of the sequences from described species came from multicellular or aggregate-forming bacteria (128 of 156, Fig. 3, SI Dataset S3). 34 bacteria encoded multiple clade 5 RTs, accounting for 30% of all the RT sequences we examined (78 of 257 clade 5 RT genes, SI Appendix Table S1). Organisms with multiple DGRs are exclusively bacteria with multicellular lifestyles, with the exception of the highly polyploid giant bacterium *Achromatium* and members of the Candidate Phyla Radiation (CPR), whose morphology remains undetermined (SI Appendix Table S1). While most of the CPR DGRs belong to the RT clade 2 (*sensu* (20), the CPR RTs in clade 5 form a distant subclade (subclade 5F, Fig. 3) and are close to the viral-encoded clade 6 RTs (20). Cyanobacteria RTs also form a previously described monophyletic subclade (subclades 5E, SI Appendix Table S1, Fig. 3) (30), which we found was almost exclusively multicellular (98%, 63 of 64 characterized species). CPR and multicellular cyanobacteria account for approximately half of the organisms with more than one clade 5 DGR RT (16 of 34), and within these two subclades, the RTs were typically closely related (SI Appendix Table S1). In contrast, multicellular *Proteobacteria, Chlorobi* and *Verrucomicrobiota* encoded more divergent clade 5 DGR RTs (Fig. 3, SI Appendix Table S1).

**Figure 3.**
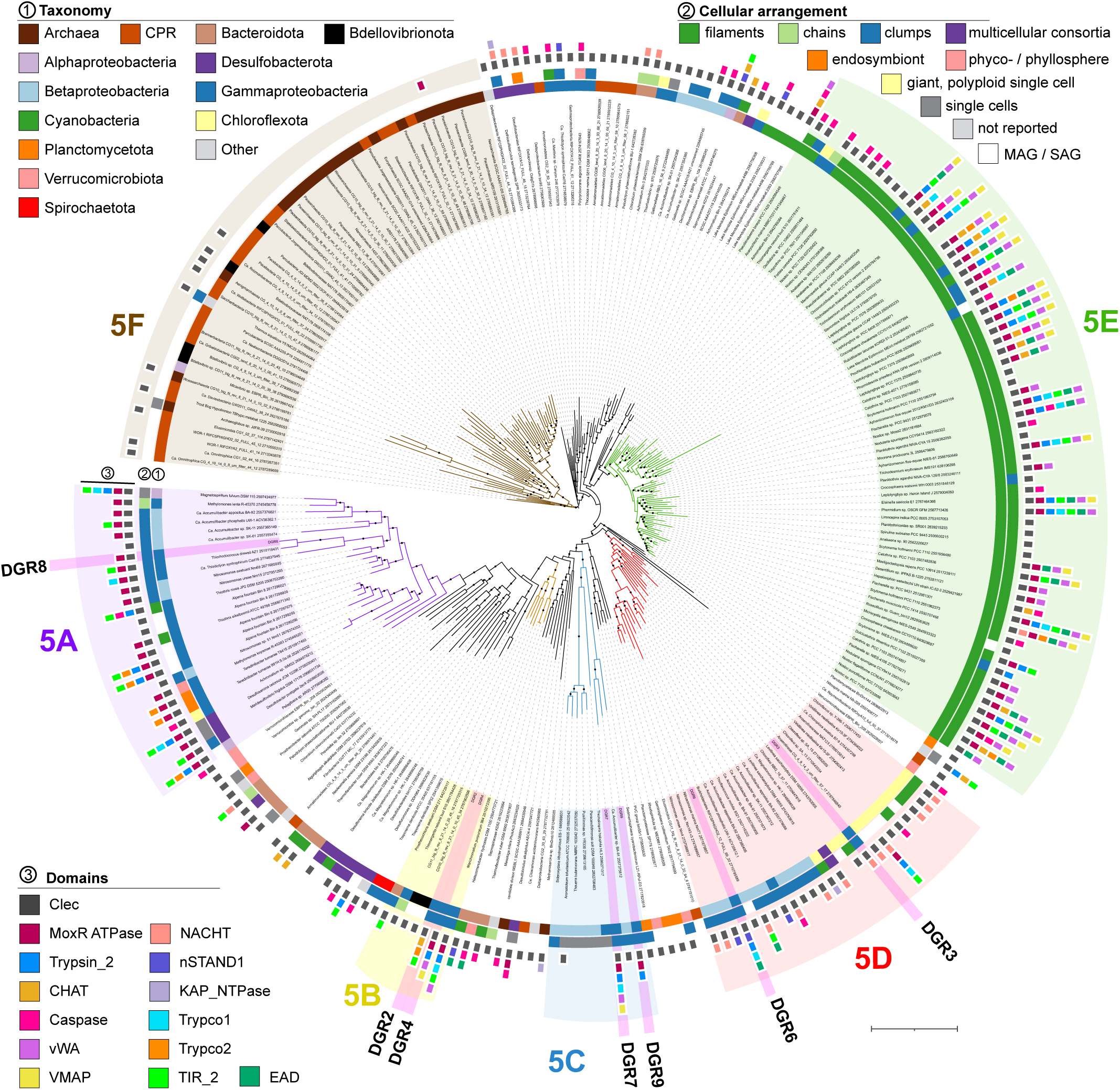
DGR reverse transcriptase (RT) genes from *Thiohalocapsa* PB-PSB1 are monophyletic with RTs from other multicellular bacteria and syntenic with conflict system domains associated with programmed cell death. The maximum likelihood phylogeny includes all clade 5 RT genes (*sensu* (20)) from genomes included in the IMG Genomes database (20), and three prior DGR surveys (19, 24, 77). The three layers of annotations indicate (from the inside to the outside): (1) the organism taxonomy, (2) the type of cellular arrangement and (3) the domains of interest detected within 20kb of the RT. Domains abbreviations and hits in each organism are provided in SI Dataset S6. The branches shaded in pink correspond to PB-PSB1’s DGR RTs. Bootstrap support greater than 70% is indicated with closed circles (n=1000). Scale bar: 1 substitution per site.

Subclade 5D, which includes RTs from *Thiohalocapsa* PB-PSB1’s DGR3 and DGR6 loci, was composed of entirely of multicellular bacteria and a large proportion of the organisms with multiple DGR loci (SI Appendix Table S1 and SI Dataset S3). Basal members of this clade, most closely related to DGR3, are filamentous chloroflexi (37–39). Derived members of this clade, more closely related to DGR6, are predominantly *Betaproteobacteria* including numerous *Accumulibacter* species from freshwater wastewater treatment reactors and marine *Nitrosomonas* that form dense microcolonies in biofiltration biofilms (40–42). Other members include the mat-forming purple sulfur bacterium *Thioflavicoccus mobilis* and a multicellular magnetotactic bacterial species from the *Deltaproteobacteria* (*Candidatus* Magnetomorum HK-1) (43, 44).

In several cases, we found organisms encoding both clade 5D and clade 5A RTs, like *Thiohalocapsa* PB-PSB1 (Fig. 3, SI Appendix Table S1). 77% of the Clade 5A sequences from described species were from multicellular organisms. This clade, like 5D, includes mostly beta*-* and gammaproteobacterial species from the *Accumulibacter*, *Nitrosomonas* and purple sulfur bacteria. PB-PSB1’s clade 5A RT (DGR8) is most closely related to the mat-forming *Thiorhodococcus drewsii*, which was isolated from the neighboring Great Sippewissett salt marsh (45) and the purple sulfur bacterium *Ca.* Thiodicyton syntrophicum, which, like PB-PSB1, forms multicellular consortia with a sulfate reducing symbiont (46).

In Clade 5B, the relatives of the DGR2 and DGR4 RTs are found in the genomes of known or putative sulfur oxidizing bacteria and, of those that have been visually characterized, all are either filamentous or aggregate forming (Fig. 3, (47–52)). The closest cultured relative to the DGR 2-4 RTs is the purple sulfur bacterium *Marichromatium purpuratum* (also from the family *Chromatiales*). Other close relatives from metagenomic data include a 40 kb *Chromatiales-*like contig from a meromictic lake (Lake La Cruz, Spain).

PB-PSB1’s DGR 7 and 9 RTs belonged to clade 5C and had few close relatives amongst cultured or high-quality MAGs in the IMG genomes database, except for *Ca*. Accumulibacter phosphatis BA-91, which also encodes a type 5D DGR (Fig. 3). Close RT relatives from unbinned metagenomic contigs (20) came from other aquatic and wastewater treatment habitats. Unlike PB-PSB1 and *Ca*. Accumulibacter phosphatis BA-91, the DGRs from the unicellular members of clade 5C were located within predicted prophage regions and targeted genes without CLec domains (or, often, any known domains, Fig. 3).

Overall, we find that one of the major cellular lineages of DGRs is composed predominantly of multicellular bacteria, many of which contain multiple distinct DGR loci like our model organism, *Thiohalocapsa* sp. PB-PSB1. PB-PSB1’s RTs belonged to distinct subclades (5A-5D) and shared similarity with diverse bacteria, illustrating their distant relationships and indicating multiple, distinct acquisitions of DGR loci by this organism (Fig. 3).

### Transposons shape the evolution of *Thiohalocapsa* PB-PSB1 DGR loci

*Thiohalocapsa* PB-PSB1’s DGRs target variable regions (VRs) in the Clec domains of 15 different genes with diverse N-terminal domains (Fig. 2). These CLec domains are quite divergent, though target genes from same DGR class typically had closely related CLec domains (SI Appendix Fig. S2). Regions of high nucleotide identity, both between and within loci, indicate that recent intragenomic duplications mediated the expansion of this organism’s DGR repertoire (Fig. 4, SI Appendix Fig. S3). Despite these footprints of duplication events, the target proteins themselves are modular, even within a class, and have highly diversified N-terminal regions encoding domains that are either completely distinct (e.g. DGR 2) or divergent homologs (e.g. the vWA domains of DGRs 4 and 5; Fig. 4, SI Appendix Fig. S3).

**Figure 4.**
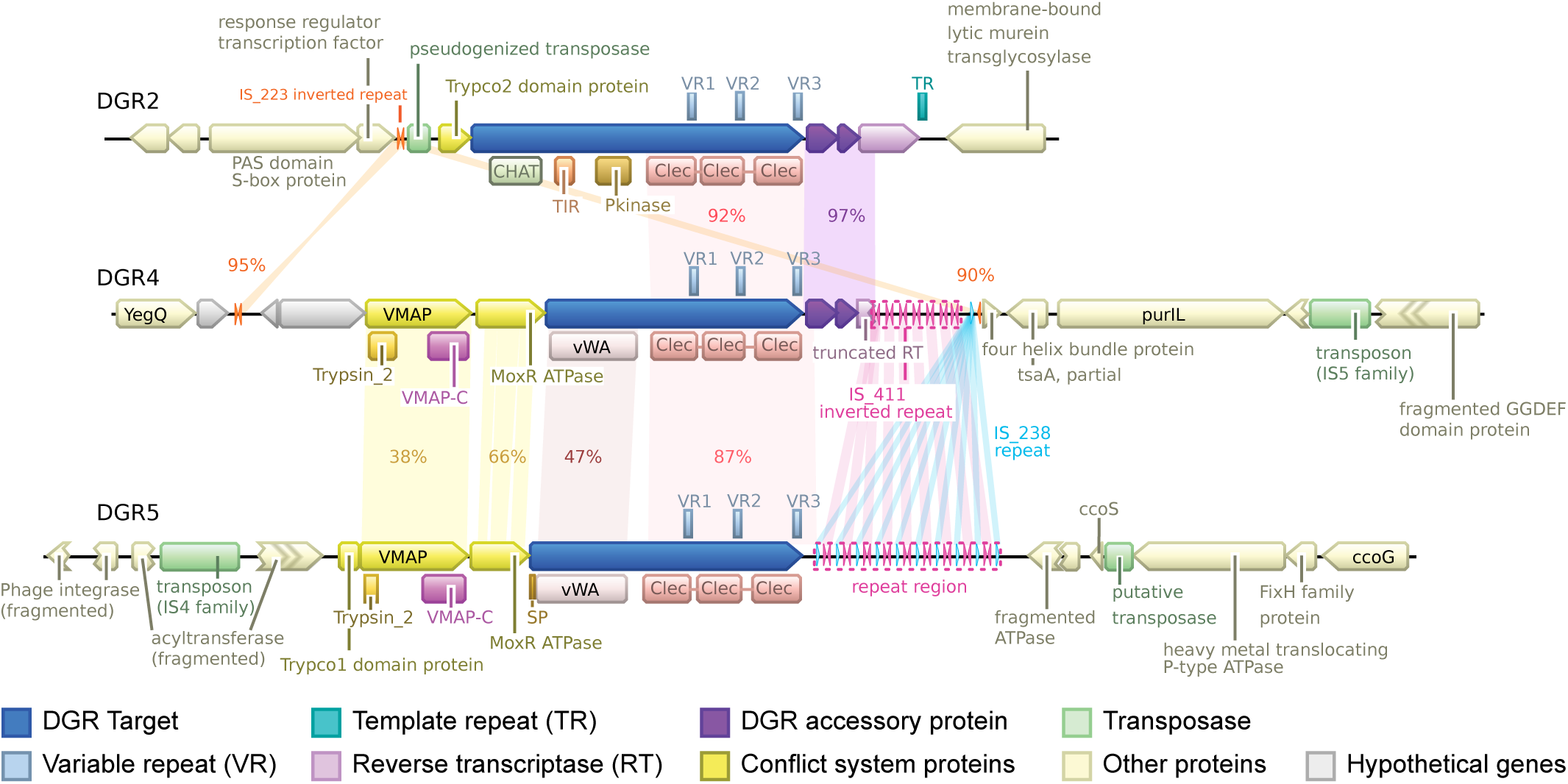
The DGR2-4-5 loci show signs of recent duplication and transposase activity. The gene neighborhoods surrounding each DGR locus are shown with regions of similarity highlighted along with the percent nucleotide identity. Regions with short direct or inverted repeats (MITE-like) are also highlighted, along with the name of the remote IS elements with matching terminal inverted repeats (IS_223, IS411 and IS_238).

The abundant insertion sequence (IS) elements of *Thiohalocapsa* PB-PSB1 (36) appear to be responsible for the duplication and divergence of DGR loci. The DGR loci contain not only complete transposons and pseudogenized transposase genes, but also miniature inverted repeat transposable element (MITE)-like sequences (Fig. 4, SI Appendix Fig. S3, SI Dataset S4). These MITE-like sequences are predicted to form stable stem loop RNA secondary structures (53) and, in some cases, matched the terminal inverted repeats from intact IS elements elsewhere in the PB-PSB1 genome. Arrays of MITE-like repeats have replaced some key functional DGR components, such as the reverse transcriptase gene at DGR 4 and 5 (Fig. 4) or the template repeat of DGR 7 (SI Appendix Fig. S3C).

The IS elements and MITE-like sequences at these loci are dynamic. Within single pink berry aggregates sequenced deeply with accurate long-read technology (PacBio HiFi), we identified structural variants with intact components alongside variants with transposon-mediated degradation (SI Appendix Fig. S4). At DGR3, we observed strains that had an intact RT gene coexisting with variants where it was interrupted by a transposon (as in the reference genome assembly). DGR 7 included strains where the RT gene and MITE-like array were deleted but also rarer variants that contained both the intact RT and a complete TR sequence (SI Appendix Fig. S4).

These examples show the prominent role mobile elements play in the evolution of DGR loci and raise the question of whether these DGRs are actively diversified or are inactive remnants of past genomic rearrangements.

### *Thiohalocapsa* PB-PSB1 DGRs are diversified in natural settings

To explore whether PB-PSB1 DGRs have diversified, we analyzed sequence variation in metagenomic data from single pink berry aggregates. Given the complexity of PB-PSB1’s DGR architecture, which has multiple duplicated regions, we started by looking for variation in the long-read sequencing (PacBio HiFi) of three individual aggregates (Fig. 5, SI Dataset S5). This analysis revealed within-aggregate variability at the expected target positions of VRs in all DGR loci except for DGR1, with different levels of variability depending on the locus and on the aggregate, indicating recent diversification of these DGR loci (Fig. 5 and SI Appendix Fig. S5). In addition to analyzing nucleotide frequencies, we summarized the variation across DGR repeats using two metrics: i) the proportion of non-reference alleles at a given position, and ii) the nucleotide diversity π within an individual aggregate, i.e. the probability that two reads from that aggregate have a different base. These metrics allow us to capture diversification at distinct scales: spatiotemporal variation from the reference genome (sampled near location A in 2011) or variation within an individual aggregate (π), which could be the sign of either recent DGR activity during the clonal growth of the colony, or of the aggregation of individual strains with distinct VR variants.

**Figure 5.**
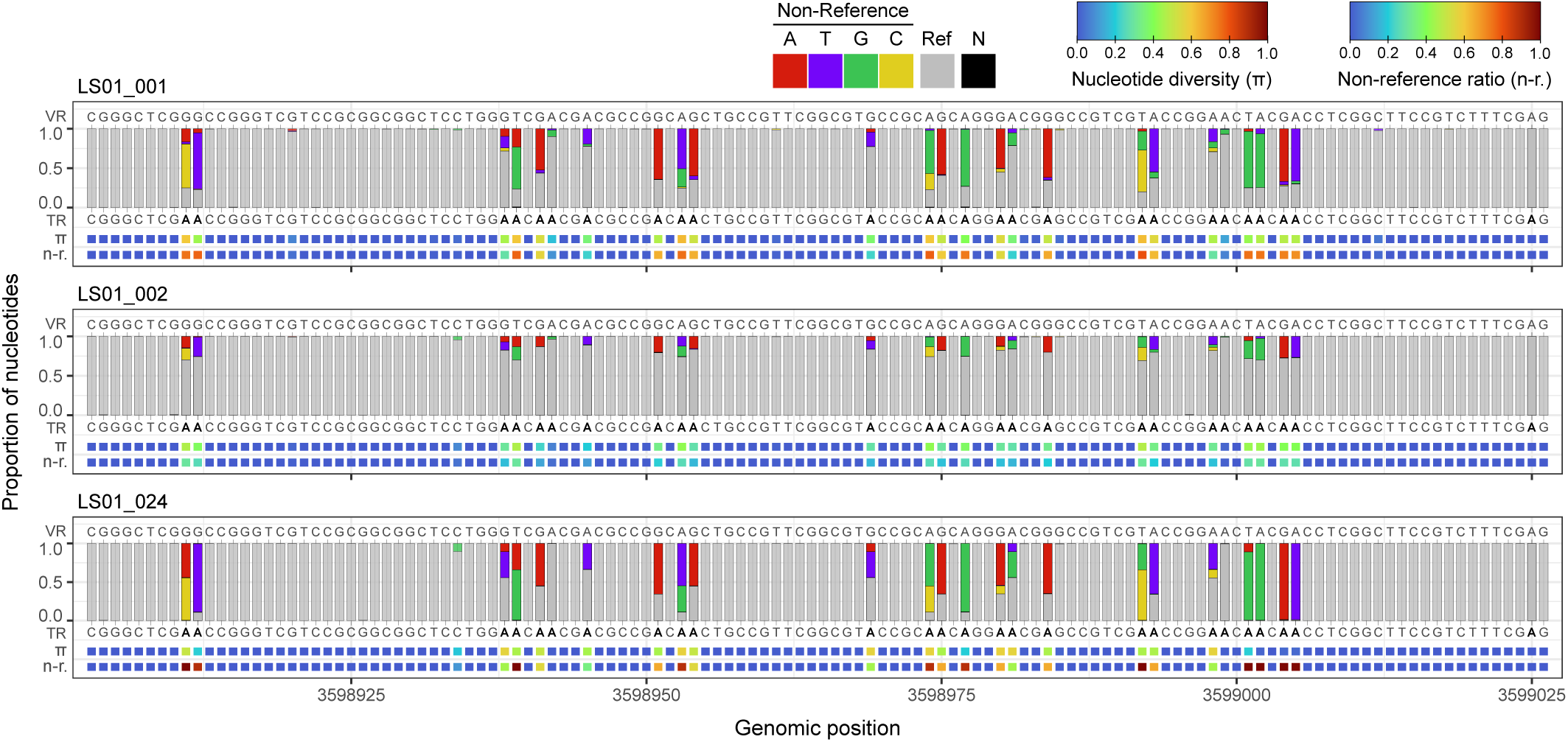
*In situ* diversification of the VR1 of DGR4 target in three pink berry aggregates from long-read metagenomic data. Each panel corresponds to an independently sequenced aggregate. Bar plots indicate the proportion of A, T, C and G at each position, colored if they differ from the reference. Letters above bars indicate the VR sequence in the reference genome, while letters below bars indicate the reference sequence of the TR with As highlighted in bold. Bottom rows show the nucleotide diversity (π) and proportion of non-reference alleles (n-r.) at each position. Ref.: Reference nucleotide. N: unknown nucleotide.

We then extended our analysis to a higher number of aggregates over the spatial range of pink berries, by searching for variation in 184 short-read metagenomes (Illumina) from single aggregates collected from five sites across three salt marshes (Fig. 1C, SI Dataset S5). Consistent with the analysis of long-read data, both the proportion of non-reference alleles and the nucleotide diversity metrics reveal signs of DGR activity in all loci except DGR1 for some aggregates in at least one sampling site (Fig. 6).

**Figure 6.**
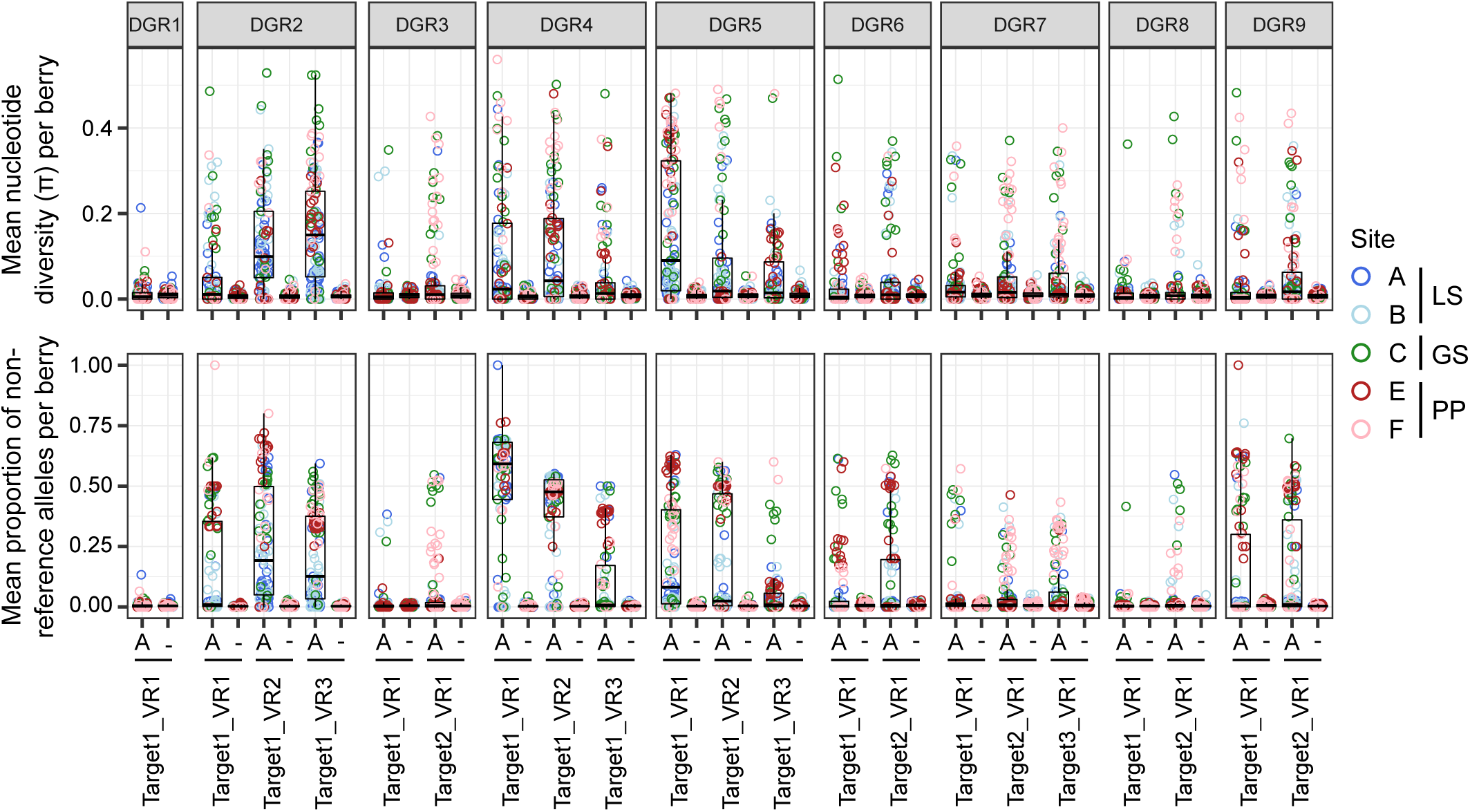
Metagenomics data reveals the differential diversification of PB-PSB1 DGR targets in 184 pink berry aggregates. For each DGR VR, the mean nucleotide diversity (upper panel) and the mean proportion of non-reference alleles (lower panel) within an aggregate were calculated separately for positions corresponding to an A in the TR (and thus targeted by mutagenesis, indicated by an A) and for all other positions (indicated by -). Each dot corresponds to a single aggregate colored by the sampling site, blue shades corresponding to Little Sippewissett (LS), green to Great Sippewissett (GS) and red shades to Penzance Point (PP). The boxplots summarize the distribution of values for all aggregates having enough coverage at a given VR.

The diversity observed at loci 3, 4, 5 and 7 was initially surprising, as they all lack key functional components in our reference genome (e.g. a complete RT or TR, Fig. 2). Thus, we explored the possibility that the targets of these loci are diversified in *trans* by an RT and/or TR encoded elsewhere in the genome. The VRs of the DGR4 and 5 targets perfectly match the TR from DGR2 at all non-adenine sites (SI Appendix Fig. S1A), suggesting that they are being diversified by DGR2 components acting in *trans*. On the contrary, the diversity observed in the targets of DGR7 cannot be explained by *trans* activity, as their VRs have no perfect match elsewhere in the genome. As mentioned above, the long read data revealed structural variants at the DGR7 locus with intact RT and TR components (SI Appendix Fig. S4). The VR diversity observed at DGR7 VRs can thus be mediated by the fraction of the population still possessing the RT and TR, and/or this diversity was generated before DGR7’s disruption by mobile elements. The structural variant with an intact VR observed at the DGR3 locus could also explain the diversity of its VRs, though as this locus possesses its own TR, it could also potentially use another RT in *trans*.

Despite the lack of variability of DGR1 target, its VR matches the DGR3 TR at all non-adenine sites (SI Appendix Fig. S1C), suggesting that DGR1 was able to mobilize DGR3’s RT and TR in *trans* before its diversification stopped.

#### Spatial patterns of DGR diversification

We observed spatial patterns in the prevalence of DGR diversification (Fig. 7). While some targets (e.g. DGR2) were diversified in most aggregates across all geographic sites, others like DGR3 were diversified frequently at some sites (C, F) but only rarely at others (A, B and E) (Fig. 7). Overall, there was a strong difference between Little Sippewissett sites A and B, where only DGR2 and to a lesser extent DGRs 4 and 5 were frequently diversified, as compared to sites in Great Sippewissett (C) and Penzance Point (E and F) where the other DGR loci were more often also frequently diversified.

**Figure 7.**
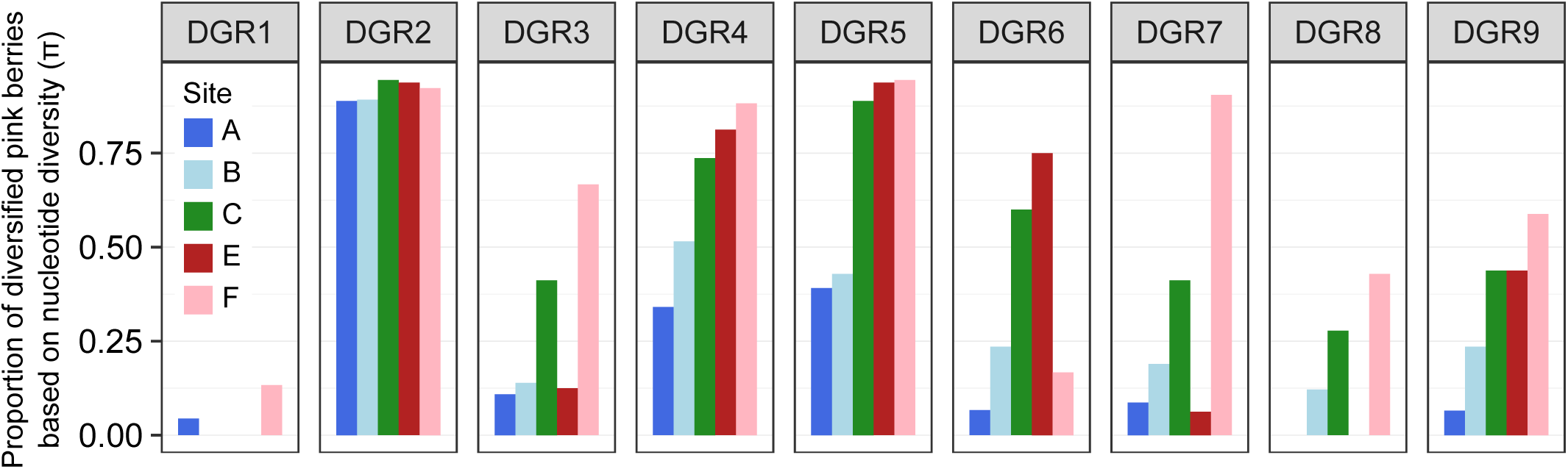
Spatial patterns in DGR diversification. The proportion of aggregates showing diversification is shown for each DGR locus at each sampling site. A DGR locus was considered to be diversified if at least one of its VRs showed diversification at positions targeted by the DGR mechanism based on the nucleotide diversity. Similar results were obtained when using the proportion of non-reference alleles. Colors correspond to sampling sites, with blue shades corresponding to Little Sippewissett (LS), green to Great Sippewissett (GS) and red shades to Penzance Point (PP). SI Appendix Fig. S6 and S7 show these results at the target and VR level, as well as the number of diversified aggregates in each case.

#### Differential activity of VRs within a DGR locus

Some PB-PSB1 DGR loci have multiple targets while others have DGR targets with multiple VR-containing CLec domains. Our metagenomic analysis revealed differences in the level of diversification of distinct targets within a given DGR locus, and of distinct VRs within a given target. For example, DGR6 has two targets, and target 1 was diversified less frequently than target 2 at sites A, B, C and F (SI Appendix Fig. S6). Similarly, for the three VRs in DGR4’s single target gene, VR3 shows variability in fewer berries than VR1 and VR2 at sites A, B and C (SI Appendix Fig. S7). This indicates that even when a corresponding RT and TR are expressed, this does not necessarily lead to equal diversification of all VRs.

Overall, the analysis of this metagenomic dataset revealed that eight of the nine DGR loci in PB-PSB1 are diversified in natural conditions, and that the level of diversity of each VR might be dependent on local environmental conditions. The expansion of active DGRs in PB-PSB1 suggests that targeted diversification is highly beneficial to this organism, and the spatial variation in activity levels raises the question of the DGR functions and triggers in this organism and in the other bacteria that possess related DGR RTs.

### *Thiohalocapsa* PB-PSB1 and other multicellular bacteria use DGRs to diversify putative antigen sensors

Several of the genes diversified by PB-PSB1’s DGRs have been identified as components of putative biological conflict systems that are enriched in multicellular bacteria (8, 13). We investigated the extent to which all of the PB-PSB1 DGRs had this type of association by annotating their neighboring genes and explored this association in the bacteria encoding the closest relatives of PB-PSB1 DGRs.

#### STAND NTPase antiviral defense systems

PB-PSB1’s DGR 3 and 6 diversify target genes with STAND family NTPase domains, as do many of the other DGRs from clade 5D (Fig. 3). STAND family NTPase domains form the central component of animal and plant innate immune responses as nucleotide-binding oligomerization-like receptors (NLRs). These sensors of pathogen-associated molecular patterns mediate programmed cell death via NLRs in large multiprotein complexes, such as the animal apoptosome and plant resistome complexes (51, 54, 55). Recent work demonstrates that bacterial STAND homologs (antiviral STAND (Avs) and bacterial NACHTs) provide protection against DNA and RNA phages *via* recognition of conserved viral proteins followed by cell death (10, 14). These experimentally characterized antiviral STAND proteins have diverse, highly variable C-terminal sensor regions, though experimental work has yet to examine STAND proteins with CLec domains.

The DGR6 architecture was described as a putative conflict system (13), with its two targets containing a STAND NTPase domain (nSTAND1) and an Effector Associated Domain (EAD8), in addition to a C-terminal formylglycine-generating enzyme (FGE) domains, a subtype of CLec folds (Fig. 8). The same EAD8 domain is also found in a nearby protein with a trypsin-like peptidase domain. The CLec domain containing the VR was proposed to be involved in sensing an invasion (13). Characterized antiviral STAND NTPases form oligomeric complexes upon sensing phage via their C-terminal region (e.g. tetratricopeptide repeats) and mediate cell death via the nuclease activity of fused N-terminal effector domains (10). Here, EAD8 domains are thought to facilitate the association of the STAND NTPase/CLec sensor and its trypsin-like protease effector by homotypic interactions (8, 13). The related DGR3 locus has a similar domain architecture, with duplicated targets and a central STAND NTPase domain (NACHT) fused to the DGR-targeted CLec domain (SI Appendix Fig. S8). While the targets at the DGR3 locus do not contain an identified adapter domain, the neighboring S8 family peptidase (N838_12175) has C-terminal tandem WD40 repeats (β-propeller). In Eukaryotes, WD40 β-propellers act as key mediators of protein-protein interactions by serving as scaffolds for multimeric assemblies that mediate diverse functions from cell division to signal transduction and apoptosis (56–58).

**Figure 8.**
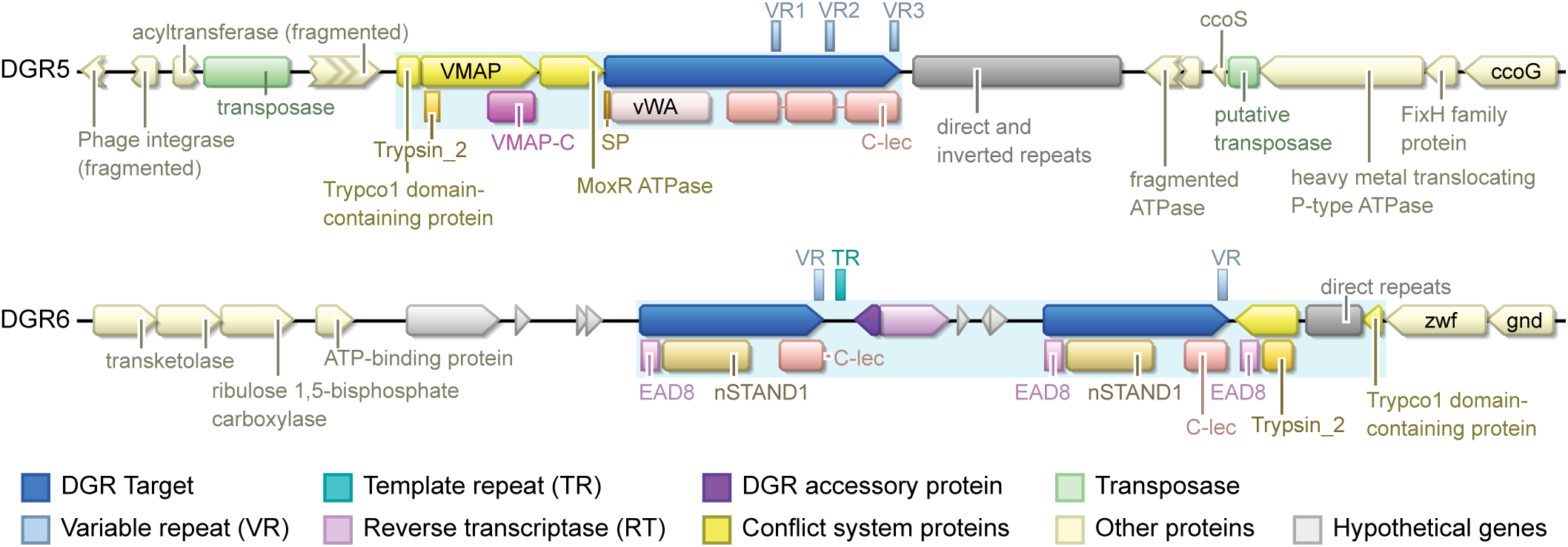
*Thiohalocapsa* PB-PSB1’s DGRs diversify putative antigen sensors in STAND NTPase and ternary conflict systems. Functional annotation of genes surrounding example ternary conflict system type (DGR5) and STAND NTPase conflict system type (DGR6) are shown. The light blue shaded areas indicate the putative DGR-containing conflict systems. SP: signal peptide, TM: transmembrane domain. The functional annotation of all DGR loci is reported in SI Appendix Fig. S8.

Like DGRs 3 and 6 from *Thiohalocapsa* PB-PSB1, DGRs from other bacteria in Clade 5D commonly had two target genes with single CLec domains and central STAND NTPase domains (nSTAND1 or NACHT), though some had other NTPases such as KAP NTPase and P-loop NTPases (Fig. 3, SI Appendix Fig. S9). Many of these targets – like DGR3 – contained N-terminal extensions without previously characterized domains, though some had known N-terminal effectors such as peptidase or TIR domains, which are key components in defense systems and cyclic-oligonucleotide based antiphage systems (7, 8, 11, 59–61). The RT subclade 5D thus seems to be specifically diversifying STAND NTPases-containing antigen sensors.

#### MoxR-like AAA ATPase ternary conflict systems

*Thiohalocapsa* PB-PSB1’s DGR 7 locus was partially described in a recent analysis of putative NTP-dependent conflict systems and has an architecture typical of the “VMAP ternary system” with a MoxR-like ATPase, a vWA domain protein, and a vWA-MoxR-associated protein (VMAP) (SI Appendix Fig. S8; (8)). Kaur *et al.* proposed that the VMAP proteins, which typically contain either a peptidase, cyclic nucleotide generating domain, or EAD, act as sensors that relay the signal of invasion to the MoxR ATPase, which activates the VMAP peptidase that in turn cleaves the vWA-domain protein liberating its associated effector domains. We propose an alternate mechanism for these ternary systems, where the DGR target protein containing both a vWA- and a variable CLec domain serves as the sensor, which relays the invasion signal through the MoxR ATPase to activate the VMAP effector, either directly or through a signal cascade.

At the DGR 7 locus, the VMAP protein has an N-terminal trypsin-like peptidase domain followed by the conserved VMAP C-terminal domain. The vWA-containing DGR target gene has a 3’,5’-cyclic AMP phosphodiesterase domain and a NACHT domain in addition to the VR-containing CLec domain, and the two additional DGR targets share the same domains except for the vWA domain (SI Appendix Fig. S8). The DGR9 locus, which belongs to the same RT phylogenetic clade (clade 5C, Fig. 3), has a relatively similar architecture, with a MoxR-ATPase directly upstream of the DGR target preceded by a trypsin-containing gene, though the DGR target contains an unidentified N-terminal extension in place of the vWA domain and the VMAP-C domain could not be detected (SI Appendix Fig. S8). None of the few cultured representatives in clade 5C had this architecture (Fig. 3). However, closer relatives to the DGR7 RT from unbinned metagenomic contigs (20) found in aquatic and wastewater treatment habitats had features of VMAP ternary systems including a MoxR ATPase and N-terminal vWA domains in the target genes (SI Appendix Fig. S9). The closest relative, a *Chromatiales-*like metagenomic contig from meromictic Lake La Cruz, had an architecture strikingly similar to *Thiohalocapsa* PB-PSB1’s DGR7, with three DGR targets containing a 3’,5’-cyclic AMP phosphodiesterase domain and a NACHT domain (SI Appendix Fig. S9). Of note, most of clade 5E DGR RTs in the multicellular cyanobacteria are associated with a MoxR ATPase and a vWA domain-containing protein, though only a subset possesses a conserved VMAP domain (Fig. 3). DGRs of this clade were speculated to have a role in signal transduction and cell death (30).

The single-target DGR4 and DGR5 loci also display the typical architecture of the VMAP ternary system. At these loci, the vWA domain is fused to the N-terminal region of the DGR target, followed by three tandem CLec domains with no other identified effector domains. Among the closest relatives of DGR4 in the RT phylogenetic tree (Subclade 5B, Fig. 3), only *Marichromatium purpuratum* 984 and an unbinned *Chromatiales* contig from Lake La Cruz possess a similar architecture of a single target gene with tandem VR-containing CLec domains that are part of a VMAP system. These target genes are syntenic with distinct vWA-domain containing genes, a MoxR-type ATPase, a peptidase domain gene (Trypsin_2, PF13365), and adapter domains (TIR_2, trypco1) as seen at the DGR 4 and 5 loci (Fig. 8, SI Appendix Fig. S8 and S9). In the other loci of clade 5B, including PB-PSB1 DGR2, there is no identified association of the DGR with a complete VMAP system. The DGR2 locus contains a Trypco2 domain that has been described as an EAD in conflict systems utilizing a trypsin-like domain as effector (SI Appendix Fig. S8) (8). The DGR2 target itself contains a caspase-like CHAT proteolytic domain and a TIR domain, which are key defense systems components (7, 8, 11, 59–61).

The DGR8 locus represents a variation on this ‘ternary’ system that was commonly associated with the large clade 5A of RTs (Fig. 3). DGR8 and other members of clade 5A have a MoxR-like ATPase immediately upstream from a target gene but in place of a vWA-domain, their target genes have N-terminal extensions without predicted domains (SI Appendix Fig. S10). DGR8 and its closest relative *T. drewsii* have a helix-turn-helix domain gene in place of the classic ternary system VMAP, suggesting an alternate mechanism to induce a cellular response (SI Appendix Fig. S9). Most of the other loci in this clade 5A had a similar organization to *Thiohalocapsa*’s DGR8, including two targets with C-terminal CLec domains, one short CLec-only target and another with a long N-terminal extension (SI Appendix Fig. S9). Clade 5A loci commonly had Trypco1, Trypco2, and TIR domains in the direct neighborhood, and often a gene containing a peptidase-like domain (either trypsin-like or caspase-like) (Fig. 3). This association of domains is interestingly similar to the one observed in the abovementioned DGR loci of multicellular cyanobacteria whose RT is quite distantly related (Clade 5E, Fig. 3).

#### Predicted cellular localization

In the PB-PSB1 genome, the target proteins in these various conflict systems are predicted to have different cellular localization (Fig. 8 and SI Appendix Fig. S8). Amongst the MoxR ATPase associated DGRs, the targets of DGR 2, 4, 7, and 8 are predicted to be cytoplasmic, while DGR5’s target is likely secreted or on the external surface, as it has a signal peptide but no transmembrane domain. DGR9’s targets contain multiple transmembrane regions that would position the CLec domains in either the cytoplasm (target 2) or the periplasm (target 1). Similarly, in PB-PSB1’s STAND NTPase systems, DGR 1 and 6 targets are likely cytoplasmic, while the NACHT and CLec domains of the DGR3 locus are predicted to be cytoplasmic in the first target and periplasmic in the second target. If the variable CLec domain is a sensor as we predict, *Thiohalocapsa* PB-PSB1 would be poised to detect threats in both the cytoplasm and periplasm.

## Discussion

### The exceptional diversification potential of PB-PSB1

Our analysis of the *Thiohalocapsa* PB-PSB1 genome revealed the presence of fourteen DGR targets that are diversified in the natural environment. Though most genomes with DGRs diversify a single target gene, genomes with multiple targets, including remote targets, are not uncommon (17, 24, 28, 30, 32, 62–64). What makes the PB-PSB1 genome exceptional is the presence of multiple DGR systems that seem to work independently. Most complete genomes that have been analyzed have no DGRs (∼98%, (20, 29)), and of the genomes with DGRs, ∼12% have two and only 2% have more than two DGRs (20). Nevertheless, our exploration of the clade 5 DGR-RT tree revealed that possessing multiple DGR loci could be relatively frequent amongst multicellular bacteria (SI Appendix Table S1), despite being exceptional in other prokaryotes. The archaeal DPANN superphylum and the ultrasmall bacteria of the Candidate Phyla Radiation were also reported to have such an expansion of DGR loci, often with multiple independent loci in a genome (19). Interestingly, the most commonly observed architecture of DGR target domains in these organisms was a CLec domain fused to an AAA ATPase (19), potentially indicating a similar mode of action to the one described here. The RT sequences of these organisms, however, form independent monophyletic clades (distant from clade 5), and their DGRs have been proposed to mediate their interaction with host organisms (19). In contrast, we demonstrate here that a specific lineage of DGRs can be recruited by diverse taxa to diversify the putative sensor protein of programmed cell death conflict systems (8, 10, 13, 14).

In PB-PSB1, the expansion of DGRs brought an unprecedented capacity for diversification, with 10^294^ possible protein combinations. As was noted in previous studies, we also observed that the positioning of the adenines in the TR is tightly constrained both to maximize diversification and avoid stop codons in the target protein (SI Appendix Fig. S1, SI Dataset S2) (16, 22, 62, 65). This highlights the strong coevolution between the target genes and their cognate TR (66). Indeed, to ensure that the DGR target gene stays functional while being diversified, selection must act on TR sequences to maintain several features: 1) An A at 1^st^ and 2^nd^ positions of codons that can be diversified without compromising the target protein integrity, 2) non-A and non-G nucleotides at the 3^rd^ position of targeted codons to avoid nonsense codons, and 3) non-As for codons that are essential and cannot be diversified. Arguably, these features could be used to recognize DGR systems under active selection.

The fact that the mutation process in DGR is not fully random but directed to the first and second positions of target codons to maximize protein diversity also explains the enrichment in non-synonymous variants in VRs reported from metagenomics analyses (20). As a consequence of this non-randomness, classical measures of selection based on dN/dS or pN/pS cannot be used to estimate the strength of the selection affecting DGR targets.

### The role of transposons in the expansion of DGRs in PB-PSB1

The PB-PSB1 DGR systems seem to have been acquired separately from distant organisms, as indicated by the RT phylogeny (Fig. 3). The dispersal of DGRs between phylogenetically distant bacteria by horizontal gene transfer (HGT) could be common, as it has been identified in several studies that pointed out the role of phages, plasmids or transposons as putative vectors (17, 19, 24, 28, 30, 63, 65). Transposons in particular have been shown to be associated with DGRs in numerous CPR bacteria and DPANN archaea (19) and appear to have induced the mobilization of a DGR to a plasmid in *Shewanella baltica*, facilitating HGT (63). In PB-PSB1, we clearly demonstrated the role of transposons in the degeneration of some DGR loci, as has been noted in other bacteria as well (63). These transposons could also be responsible for DGR loci duplications either by translocating parts of DGRs as a cargo, or by triggering homologous recombination between their numerous copies distributed within the genome. Several systems seem to have diversified within PB-PSB1 through intragenomic duplication and/or recombination of either a target (e.g. the two targets of DGR6) or a partial system (e.g. DGR2-4-5). The latter example is particularly striking: the RT, accessory genes and the target gene C-terminal region have been duplicated and recombined with distinct N-terminal domains, while MITE-like tandem repeats have degraded redundant DGR components (Fig. 4). More generally, duplication/recombination has been proposed as a mechanism to extend the repertoire of DGR target proteins by adding VR-containing CLec domains to the N-terminal end of diverse proteins (20, 26, 30, 63). DGR systems thus combine two different dimensions of variation, domain shuffling and hypermutation, which are particularly well exemplified in PB-PSB1.

### Diversification is tightly constrained by the local environment

The spatial patterns in the diversity of PB-PSB1’s predicted conflict systems likely reflect differences in the biotic pressures acting on this organism throughout its geographic range. While a prior analysis of metagenomic time series data showed that most DGRs were used to maintain population-level diversity in the DGR target sequences (20), our metagenomic data reveals that this diversity is spatially constrained for PB-PSB1. Indeed, the level of diversification of each DGR target, and even of each VR within a target gene, depends on the location of the pink berry aggregates across salt marshes less than 10 km apart. This spatial pattern of DGR diversification could result from 1) differential activity of DGR regions depending on local environmental conditions, with specific conditions triggering the activation of each DGR, 2) different selective pressures at each site, or 3) a combination of both. At some sites, select DGR targets showed very low levels of diversity (e.g. DGR 3 and 8 in Little Sippewissett sites A, B and LS01, Fig. 7 and SI Appendix Fig. S5). This indicates either inactivity due to repression or disruption of these systems, or the presence of strong purifying selection favoring the reference allele in a particular geographic region. To discriminate between these processes, one could examine if distinct VR variants or combinations of variants were found in different locations. However, long-read metagenomes of high sequencing depth across several sites in all three salt marshes would be needed to phase single nucleotide variants. Paul and Eren (67) recently used metagenomes to compare the variation in *Trichodesmium erythraeum*’s DGR targets over a much larger spatial scale between the Pacific and Indian oceans. They showed the presence of distinct combinations of variants at these two locations, which is likely the result of differential selection. Of note, our results showed some population heterogeneity in the disruption of DGR loci, demonstrating that they are under a balance between a relaxed selection allowing for the degradation of the system, and a selection for functional, diversifying systems. Regardless of the underlying mechanism, the spatial pattern of diversification highlights the importance of exploring DGR activity in various environmental or experimental conditions (20, 67).

### DGR targets as sensors in conflict systems of multicellular bacteria

Our manual annotation of DGR targets and surrounding genes revealed that most, if not all, of the PB-PSB1 DGRs are parts of putative conflict systems where we believe the diversified CLec domains of the DGR targets are used as antigen sensors. Within each conflict system, it is likely that the proteins associate into a complex (8, 10, 13), thus increasing the avidity of the CLec domains towards specific antigens (66, 68). Amongst described antiviral STAND NTPase systems, binding of conserved viral-associated molecular patterns initiates complex formation and programmed cell death (10), an effector response conserved across animal and plant innate immune responses and also proposed for ternary conflict systems (8, 13). In PB-PSB1, the variable domains are predicted to reside in different cellular locations (cytoplasmic, extracellular, membrane associated), and would equip bacteria to sense invasions in all cellular compartments. This proposed interaction between diversifying sensors and invading antigens, however, raises the intriguing question of how the systems distinguish self *vs* non-self to avoid autoimmunity.

A striking proportion of DGRs from this major lineage (clade 5) are encoded by diverse bacteria with multicellular or aggregative lifestyles (Fig. 3, SI Dataset S3). These bacteria often had multiple DGR systems that diversify similar putative antigen sensors with lectin and NLR domains (Fig. 3). In an arms-race scenario, the benefits of hypervariable antigen sensors for the surveillance and response to novel threats are clear, particularly to bacteria with multicellular or aggregative lifestyles: the cell density and limited genetic diversity within both single and multi-species biofilms or aggregates makes them especially vulnerable to attacks by viruses or mobile DNA/RNA. The potential parallels to the vertebrate adaptive immunity are striking: (1) diversification of a pattern recognition sensor from innate immunity, (2) targeted mutation restricted to a short region of the sensor gene via error-prone DNA synthesis/repair, (3) clonal expansion of cells with high affinity sensors (33, 69).

These parallels raise interesting evolutionary questions about the emergence of multicellularity. We propose that infection represents an important and universal evolutionary constraint for all cellular aggregations of close genetic relatives. The risk of infectious epidemics in an aggregate with little genetic diversity creates a strong selective pressure for successful multicellular life forms, from bacteria to metazoans, to evolve sophisticated immune responses that include threat surveillance by highly diversified sensors, as well as programmed cell death. In eukaryotes, programmed cell death is seen as necessary for multicellular life (54), and in model simulations, the evolution of programmed cell death has been coupled to the evolution of multicellularity, as it becomes beneficial for bacterial immunity once cells aggregate (70).

In addition to a vulnerability to attacks, multicellularity means sharing resources with neighbors, introducing a selection pressure for kin recognition (71). Interestingly, the NLR domains (such as the STAND-NTPase domains of PB-PSB1 DGR3 and other subclade 5D targets) are homologous to some fungal hetero-incompatibility determinants (10, 14, 72). In these fungi, where cell fusion and syncytial organization are common, the hetero-incompatibility systems prevent fusion with genetically distinct strains by triggering localized, programmed cell death (71, 72). This suggests an intriguing alternative role for the DGR-diversified conflict systems in multicellular bacteria: to permit the association with cells sharing similar alleles and avoid associations with non-kin cells, in a similar fashion to fungi hetero-incompatibility or bacterial kin recognition systems (71, 73).

## Conclusions

Here, we used *Thiohalocapsa* PB-PSB1 as a model to study the association of DGRs with bacterial conflict systems. This revealed a genome with an unusually high number of diversity-generating retroelements, bringing an astronomical potential for targeted diversification. PB-PSB1 DGR RTs were found to belong to a monophyletic clade, which we show to contain error-prone RTs from diverse multicellular bacteria. We propose that these DGRs act as variable antigen sensors in conflict systems triggering programmed cell death, either as a defense mechanism or for kin recognition. Based on metagenomic data, we show that 14 target proteins of PB-PSB1 DGRs are diversifiedacross the bacteria’s known geographic range, with spatial patterns in the level of diversity of each locus likely due to differential biotic pressures either regulating the conflict systems or selecting specific variants. Overall, our study highlights how hypermutation of antigen sensors could represent an evolutionary response to the constraints of multicellularity, and calls for a molecular characterization of these systems in multicellular prokaryotes.

## Material and methods

Extended methods are available in the SI Appendix Supporting Text.

### Sequence data used in this study

The circular metagenome-assembled genome of *Thiohalocapsa* PB-PSB1 was downloaded from Genbank (NCBI accession number: GCA_016745215.1).

For metagenomics, 187 pink berry aggregates were sampled from 6 ponds across 3 salt marshes near Woods Hole, MA (Fig. 1, SI Dataset S5). 184 aggregates sampled between 2015 and 2017 were sequenced on an Illumina HiSeq 2500 (250bp paired-end reads) at the Whitehead Institute for Biomedical Research (Cambridge, MA). Illumina reads were cleaned with bbduk.sh in bbmap v38.92 (https://sourceforge.net/projects/bbmap/) with options forcetrimright2=30 qtrim=rl trimq=20 maq=20 minlen=50.

Three additional aggregates sampled in 2022 were sequenced for long-reads metagenomics on a Pacific Biosystem Sequel IIe sequencer. CCS reads were filtered for duplicates using pbmarkdup, and BBMap was used to remove potential chimeric reads, and trim adapters.

### Detection of DGRs in Thiohalocapsa PB-PSB1’s genome

The PB-PSB1 genome was run on the MyDGR web server (74) with default options. The DGR detection analysis described in (19) was also performed using the python package available at https://pypi.org/project/DGR-package/ (19, 20). All predicted DGR loci were manually inspected. myDGR correctly predicted the DGR7 locus based on the presence of the RT gene, but wrongly identified the target genes because of the missing TR. An alignment of CLec domains found at this locus to the DGR9 TR revealed the positions of the correct VRs.

### Annotation of transposons and MITE-like sequences

Insertion sequence (IS) elements in *Thiohalocapsa* PB-PSB1’s genome were annotated using ISEScan v1.7.1 with default parameters (75) (SI Dataset S4). Three IS elements were predicted within the target genes of the DGR7 locus, but a manual inspection revealed that these did not contain a transposase gene and were likely an overprediction by ISEScan. Regions of short direct and inverted repeats at the DGR loci were identified by manual inspection of dotplots, and their annotation was refined using Find Repeats in Geneious Prime 2022.1.1 (https://www.geneious.com). These repeats were searched against a database of terminal inverted repeats from the intact IS elements detected with ISEScan using BLASTn (-word_size 7 -gapopen 3 -gapextend 2 -reward 1 -penalty -1). Short inverted repeats matching existing IS-elements were analyzed with RNA Fold (76). Stable hairpin-forming inverted repeats were characterized as Miniature Inverted-repeat Transposable Elements (MITE)-like sequences.

### Phylogenetic analysis of DGR reverse transcriptases

The seven DGR RTs from PB-PSB1 were compared to a reference database of 1143 representative RT sequences (20) following Roux *et al.* phylogenetic pipeline, allowing for the identification of the closest relatives of PB-PSB1 DGRs in IMG genomes and metagenomes. As all PB-PSB1’s RTs fell within RT clade 5 (*sensu* (20), we performed a second phylogenetic analysis of this clade with a refined and expanded dataset (SI Dataset S3). This new dataset included all clade 5 RT genes from genomes in the IMG Genomes database (20), and three prior DGR surveys (19, 24, 77). RT sequences from metagenomes or single-cell genomes were filtered out, except those present in the IMG Genomes database. The 257 amino acid sequences were aligned with MAFFT v7.407 (78) using the einsi mode and automatically trimmed using TrimAl v1.4.rev15 (79) with the -gappyout option. The RT tree was built with IQ-Tree v1.5.5 (80) using the built-in model selection (optimal model: Q.pfam+F+R10) (81) and 1000 bootstrap replicates.

iTOL (82) was used to render the tree and visualize domains associated with the clade 5 DGR RT loci. For all members of the tree, select conflict system-associated domains within 20 kb of the DGR RT were identified either from the IMG annotations of PFAM domains or via *hmmscan* (83) (SI Dataset S6).

### Detection of protein domains in DGR-containing loci

Protein sequences of each DGR target and neighboring genes in *Thiohalocapsa* PB-PSB1 and a selection of genomes and metagenomic contigs with RTs in clades 5A-5D were manually inspected to refine their annotations. Each sequence was run through InterProScan (84), hmmscan in the HMMER Web server (85) and NCBI Conserved Domains Database (86, 87) to predict functional domains. In addition, the protein sequences were compared to the domains described in (8, 13) using *hmmscan* in HMMER 3.3.2 (83), based on PFAM hmm profiles and the vWA-ternary domain alignment provided by the authors (SI Dataset S7).

### Calculation of the number of potential protein sequence combinations

A custom python script (available at https://github.com/hdore/PB-PSB1_DGR_variation) was developed to calculate all possible protein sequence combinations for each VR of each DGR. The script uses the coordinates of the target protein and VR, and the sequence of the TR to identify the codons (and the positions within each codon) targeted by DGR. It counts all the potential amino acids that can be generated by changing nucleotides at all targeted positions of the codon. To be more accurate for DGR7 locus we used the TR sequence identified in a long-read structural variant (see Results).

### Detection of DGR activity and structural variants from metagenomics data

To detect the footprints of DGR activity from metagenomics data, the PacBio HiFi and Illumina reads were mapped to the PB-PSB1 genome using minimap2 v2.24-r1122 (option -ax map-hifi) (88) or *bwa mem* (bwa v0.7.17-r1188, default options) (89), respectively. The resulting *bam* alignment files were filtered using samtools (90) to keep only mapped reads, and Illumina reads were further filtered with a custom python script to keep only reads mapping at least partially onto a DGR VR and whose mate mapped within 1kb, preventing unspecific mapping between similar DGR targets. samtools *mpileup* was then run with option -a to get a pileup of reads at each position of the genome. Custom python scripts were used to extract the proportion of each nucleotide (A, T, C, G), at each position of each VR and to calculate the proportion of non-reference alleles and the nucleotide diversity (π = 1-(a^2^+t^2^+c^2^+g^2^) with a, c, t and g being the proportion of each nucleotide (91)) at each position. These metrics were calculated only at positions that had more than 5x coverage.

We considered a VR to be diversified in a given “pink berry” aggregate metagenome if

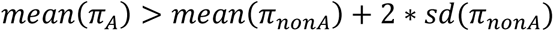

With π_*A*_the mean value of nucleotide diversity at VR positions targeted by DGR and π_*nonA*_the mean value of nucleotide diversity at VR positions *not* targeted by DGR. The same formula was applied for the proportion of non-reference alleles. When aggregating results at the target- or DGR-level, a locus was considered as diversified in an aggregate if at least one of the VRs of this locus showed variability.

The *bam* alignment files of PacBio long reads were also used to detect potential structural variants at DGR loci by manual inspection using IGV (92) and GenomeRibbon (93). In the case of DGR 7, a read representing the structural variant with an intact TR was identified in this manner and annotated using MyDGR (SI Appendix Fig. S4D, SI Dataset S7).

## Acknowledgements

We thank L. Aravind for providing sequence alignments and hmm profiles of the domains his group described as being involved in putative conflict systems. We also thank José T. Saavedra for the “pink berry” photograph shown in Fig. 1. Some figure panels were created with BioRender.com. This work was supported by a Whitman Fellowship to EGW from the Marine Biological Laboratory and long read sequencing was supported by a Joint Genome Institute New Investigator CSP award to EGW (508543). Use was made of computational facilities purchased with funds from the National Science Foundation (CNS-1725797) and administered by the Center for Scientific Computing, which is supported by the California NanoSystems Institute and the Materials Research Science and Engineering Center (MRSEC; NSF DMR 2308708) at UC Santa Barbara.

## Data sharing plans

All new sequencing data is available in NCBI GenBank under BioProject PRJNA1019683 and JGI’s IMG database (3300056627, 3300056928, 3300056818). Accession numbers for each sample are provided in SI Dataset S5. Code used in the analysis is available at https://github.com/hdore/PB-PSB1_DGR_variation.

