## Supplemental Appendix for "Targeted hypermutation of putative antigen sensors in multicellular bacteria"

**This PDF file includes:**

Supporting text
Legends for Datasets S1 to S8
Table S1
Figures S1 to S9
SI References

**Other supporting materials for this manuscript include the following:**

Datasets S1 to S9

### Supporting Information Text

### Extended Methods

#### DNA extraction and sequencing

To analyze the diversity of PB-PSB1 in its natural environment, 187 pink berry aggregates were sampled from 6 ponds across 3 salt marshes near Woods Hole, MA (Figure 1, Supplemental Data 5). 184 metagenomes from individual aggregates sampled between 2015 and 2017 were sequenced with short-read sequencing technology. DNA was extracted from all samples with the Agencourt DNAdvance Genomic DNA Isolation Kit (Beckman Coulter, Indianapolis, USA). Metagenomic libraries were prepared with the Nextera XT DNA Library Prep Kit and Illumina index primers (Illumina, San Diego, USA). Libraries were quantified on an Agilent 4200 TapeStation system with High Sensitivity D5000 ScreenTapes (Agilent Technologies, Santa Clara, USA) and pooled by equimolar amounts. Sequencing was performed on an Illumina HiSeq 2500 machine (250bp paired-end reads) at the Whitehead Institute for Biomedical Research (Cambridge, MA). The resulting Illumina reads were cleaned with bbduk.sh in bbmap v38.92 (https://sourceforge.net/projects/bbmap/) in two steps. First, the last 30bp at the 5' end of reads were removed with option forcetrimright2=30. Then read ends were trimmed based on quality (Q20) and only reads longer than 50 bp with an average quality above Q20 were kept (options qtrim=rl trimq=20 maq=20 minlen=50).

Three additional aggregates sampled in 2022 were sequenced with PacBio HiFi long-read technology. After sampling, aggregates were rinsed with 0.22 µm-filtered seawater and frozen at -80 °C until DNA extraction. In order to maintain DNA integrity for long-read sequencing, a DNA extraction protocol was adapted from (1) and (2). Briefly, a single frozen aggregate was ground in a 1.5 mL tube and incubated at 37 ºC for 1h with 125 µL of Tris Lysis Buffer with extra EDTA (100 mM NaCl, 10 mM Tris HCl at pH 8, 100 mM EDTA at pH 8, 0.5% w/v SDS) and 10 µL of lysozyme at 100 mg/mL. After addition of 125 µL of warm 4% high-salt CTAB (4% w/v CTAB, 10 mM Tris HCl pH 8, 100 mM EDTA pH 8, 2.8 M NaCl), 6 µL of proteinase K (20 mg/mL, New England Biolabs, Ipswich, MA, USA) and 0.8 µL RNAse A (New England Biolabs, Ipswich, MA, USA), the tube was incubated at 55 ºC for 3 hours. After cooling down, 250 µL of chloroform:isoamyl alcohol (24:1) were added before mixing for 15 min on a rotator mixed. The aqueous and organic phases were then separated by spinning for 15 min at 6000 g, and the aqueous phase was carefully transferred to a fresh 1.5 mL tube. This cleaning step was repeated a second time to remove any trace of proteins. Then, 450 µL of warm CTAB precipitation buffer (2% w/v CTAB, 50 mM Tris HCl pH 8, 100 mM EDTA pH 8) were added. After overnight incubation at 55 ºC, the precipitated DNA was harvested by centrifugation at 16000g for 15 min. The DNA pellet was rinsed twice with cold 80% ethanol before elution in 10 mM Tris HCl, pH 8.

To prepare PacBio HiFi libraries, an input of 50 ng of genomic DNA was sheared to 6 kb - 10 kb using the Megaruptor 3 (Diagenode). The sheared DNA was treated with an exonuclease to remove single-stranded ends, a DNA damage repair enzyme mix, and an end-repair/A-tailing mix, and then ligated with amplification adapters using SMRTbell Express Template Prep Kit 2.0 (PacBio). Templates were purified with ProNex Size-Selective Purification System (Promega). The purified ligation product was split into two reactions and enriched using 10 cycles of PCR using the SMRTbell gDNA Sample Amplification Kit (PacBio). The amplified product was combined and treated with a DNA damage repair enzyme mix and an end-repair/A-tailing mix and ligated with barcoded overhang adapters. Libraries were size-selected using the 0.75% agarose gel cassettes with Marker S1 and High Pass protocol on the BluePippin (Sage Science). The PacBio Sequencing primer was then annealed to the SMRTbell template library and sequencing polymerase was bound to them using Sequel II Binding kit 2.0. The prepared SMRTbell template libraries were sequenced on a Pacific Biosystem Sequel IIe sequencer using SMRT Link 10.2, tbd-sample dependent sequencing primer, 8M v1 SMRT cells, and Version 2.0 sequencing chemistry with 1x1800 sequencing movie run times. CCS reads were processed with the JGI QC pipeline to remove artifacts. Briefly, reads were filtered for duplicates using pbmarkdup, analyzed using the icecreamfinder.sh script in BBMap to filter potential chimeric reads, and adapter trimmed using bbduk.

**Annotation of MITE-like sequences**

Regions of short direct and inverted repeats at the DGR loci were identified by manual inspection of dotplots and their annotation was refined using Find Repeats with Geneious Prime 2022.1.1 (https://www.geneious.com). These repeats were searched against a database of terminal inverted repeats from the intact IS elements detected with ISEScan using BLASTn with parameters adjusted for short search sequence and to maximize hits covering the entire repeat length (-word_size 7 -gapopen 3 -gapextend 2 -reward 1 -penalty -1). Short inverted repeats matching existing IS-elements were analyzed with RNA Fold (http://rna.tbi.univie.ac.at/cgi-bin/RNAWebSuite/RNAfold.cgi) (3). Stable hairpin-forming inverted repeats were characterized as Miniature Inverted-repeat Transposable Elements (MITE)-like sequences.

### Legends for Datasets S1 to S8

**Dataset S1 (separate file).**
Coordinates of PB-PSB1 DGR elements. Note that when the element is on the minus strand, the end coordinate is greater than the start coordinate. When multiple VRs are present in a target, they are numbered from the N-terminal end to the C-terminal end. All VR to TR coordinates were verified manually based on VR/TR alignment.

**Dataset S2 (separate file).**
Calculation of diversification potential for each VR of each DGR target in PB-PSB1. The codons that are targeted by DGR were identified based on the TR/VR alignment (corresponding to positions were the TR has an A). Then the number of potential amino acids (or stop codons) that can be generated by DGR-induced mutation from each codon was calculated using the standard bacterial genetic code. Note that while the codons are identified on the target genes, the codon reference sequence is based on the aligned TR sequence, as the DGR mechanism replaces the whole VR with the TR sequence, including non-variable positions. The table includes the sequence of targeted codons and the positions that are targeted by DGR within each codon. The smaller table on the right-hand side summarizes the results by VR, target and DGR locus. For DGR7 the TR sequence extracted from the long read presenting a structural variant with the intact TR was used for more accuracy.

**Dataset S3 (separate file).**
Description of all clade 5 DGR-encoding organisms. The metadata available in the IMG Genome database was manually curated, and additional characteristics such as the cell morphology and the multicellular status were added based on the literature cited in the column "Morphology_citation". The IMG accession numbers of the genome and the reverse transcriptase gene are indicated.

**Dataset S4 (separate file).**
Results of ISEScan detection of insertion sequences (IS) elements. This table corresponds to the raw output from ISEScan. TIR: Terminal Inverted Repeat. Tpase: transposase.

**Dataset S5 (separate file).**
Single-aggregate metagenomes metadata and accession numbers.

**Dataset S6 (separate file).**
Annotation of a selection of conflict system-associated domains in clade 5 RTs neighborhood. The first tab indicates the domains that were searched for and their abbreviations used in Fig. 3 and in the main text. These domains were searched for within 20 kb of the DGR clade 5 RTs. For each domain, a separate tab indicates the domain hits in the organisms represented on Fig. 3. The IMG accession number of the genome, the RT gene and the gene with a domain hit are indicated.

**Dataset S7 (separate file).**
Amino-acid alignment of the vWA domains, including vWA sequences from ternary conflict systems, as used by (4) to build a hmm profile. The hmm profile was used to search for vWA domains in DGR neighborhoods.

**Dataset S8 (separate file).**
Genbank file containing the MyDGR annotation of a long read presenting a structural variant of Thiohalocapsa PB-PSB1’s DGR7 locus with an intact template repeat.

**Table S1.** Distribution and multicellular status of organisms containing more than one clade 5 error-prone reverse transcriptase gene. Cyanobacteria are shaded in green and CPR are shaded in brown. Average phylogenetic distance is the mean of pairwise phylogenetic distance for all clade 5 RT genes in a genome.

| Nb of full-length RTs | Average RT Phylogenetic Distance | Number of RTs within each clade 5 subclade | | | | | | | Species | Multicellular or aggregate-associated | IMG Genome ID |
|---|---|---|---|---|---|---|---|---|---|---|---|
| | | 5A | 5B | 5C | 5D | 5E | 5F | Other | | | |
| 5 | 3.45 | DGR8 | DGR2-4 | DGR7-9 | DGR3-6 | | | | Thiohalocapsa sp. PB-PSB1 (all RTs included | yes | 2867970272 |
| 4 | 1.79 | | | | 1 | | | 3 | Ca. Magnetomorum sp. HK-1 | yes | 2648501189 |
| 3 | 2.85 | 1 | | | 2 | | | | Ca. Accumulibacter appositus BA-92 | yes | 2556921088 |
| 3 | 2.75 | 1 | | | | | | 2 | Ca. Accumulibacter sp. SK-01 * 2 RTs @ 1 locus | yes | 2556921083 |
| 3 | 1.24 | | | | | 3 | | | Achromatium Bin 0 | no - highly polyploid | 2642422597 |
| 3 | 0.64 | | | | | 3 | | | Calothrix sp. PCC 7103 | yes | 2507262048 |
| 3 | 0.51 | | | | | 3 | | | Scytonema hofmanni PCC 7110 | yes | 2551306141 |
| 2 | 4.10 | 1 | | | | | | 1 | Ca. Thiodictyon syntrophicum Cad16 | yes | 2773857920 |
| 2 | 3.81 | 1 | | | 1 | | | | Ca. Accumulibacter phosphatis UW-1 | yes | CP001715.1 |
| 2 | 3.37 | 1 | | | 1 | | | | Verrucomicrobiaceae EBPR_Bin_208 | yes | 2619618930 |
| 2 | 3.33 | 1 | | | 1 | | | | Ca. Accumulibacter sp. SK-11 | yes | 2556921085 |
| 2 | 3.25 | | | 1 | 1 | | | | Ca. Accumulibacter sp. BA-91 | yes | 2556921087 |
| 2 | 2.74 | | | | | | 2 | | Bdellovibrionaceae NAT178 | unknown | 2802429364 |
| 2 | 2.54 | | | | | | | 2 | Pelodictyon phaeoclathratiforme BU-1 | yes | 642555146 |
| 2 | 2.21 | | | | | | | 2 | Thermoflexibacter ruber DSM 9560 | yes | 2636415974 |
| 2 | 1.93 | | | | | | | 2 | Phaeodactylibacter xiamenensis KD52 | yes | 2617271238 |
| 2 | 1.69 | | 1 | | | 1 | | | Thiomargarita nelsonii bud S10 | yes | 2600255314 |
| 2 | 1.35 | | | | | | 2 | | Parcubacteria GW2011_GWA2_45_13 | unknown | 2626541992 |
| 2 | 1.26 | | | | 2 | | | | Viridilinea mediisalina Kir15-3F | yes | 2751186036 |
| 2 | 1.06 | | | | | 2 | | | Trichodesmium erythraeum IMS101 | yes | 637000329 |
| 2 | 1.00 | | | | | 2 | | | Crocosphaera chwakensis CCY0110 | yes | 640612201 |
| 2 | 0.99 | | | | 2 | | | | Levilinea saccharolytica DSM 16555 | yes | 2740892510 |
| 2 | 0.95 | | | | | | 2 | | Ca. Woesearchaeota CG10_big_fil_rev_8_21_14_0_10_30_7 | unknown | 2785511156 |
| 2 | 0.91 | | | | | | 2 | | Ca. Staskawiczbacteria RIFOXYB1_FULL_32_11 | unknown | 2711768669 |
| 2 | 0.75 | | | | | 2 | | | Nostoc sp. PCC 7120 | yes | 637000199 |
| 2 | 0.67 | | | | | 2 | | | Merismopedia glauca CCAP 1448/3 | yes | 2802429465 |
| 2 | 0.65 | | | | | 2 | | | Lake Mendota Epilimnion MEint.metabat.2353 | unknown | 2582580551 |
| 2 | 0.63 | | | | | 2 | | | Nodularia spumigena CCY9414 | yes | 2562617131 |
| 2 | 0.63 | | | | | 2 | | | Lake Mendota Epilimnion MEint.metabat.4498 | unknown | 2582580576 |
| 2 | 0.51 | | | | | 2 | | | Fischerella sp. PCC 9431 | yes | 2512875027 |
| 2 | 0.48 | | | | | | 2 | | Ca. Pacearchaeota CG10_big_fil_rev_8_21_14_0_10_34_12 | unknown | 2785510794 |
| 2 | 0.42 | | | | | | 2 | | Armatimonadetes CG06_land_8_20_14_3_00_66_21 | unknown | 2786546208 |
| 2 | 0.35 | | | | | | 2 | | Ca. Pacearchaeota CG10_big_fil_rev_8_21_14_0_10_30_48 | unknown | 2785510795 |
| 2 | 0.33 | | | | | 2 | | | Leptolyngbya sp. PCC 7375 | yes | 2509601039 |

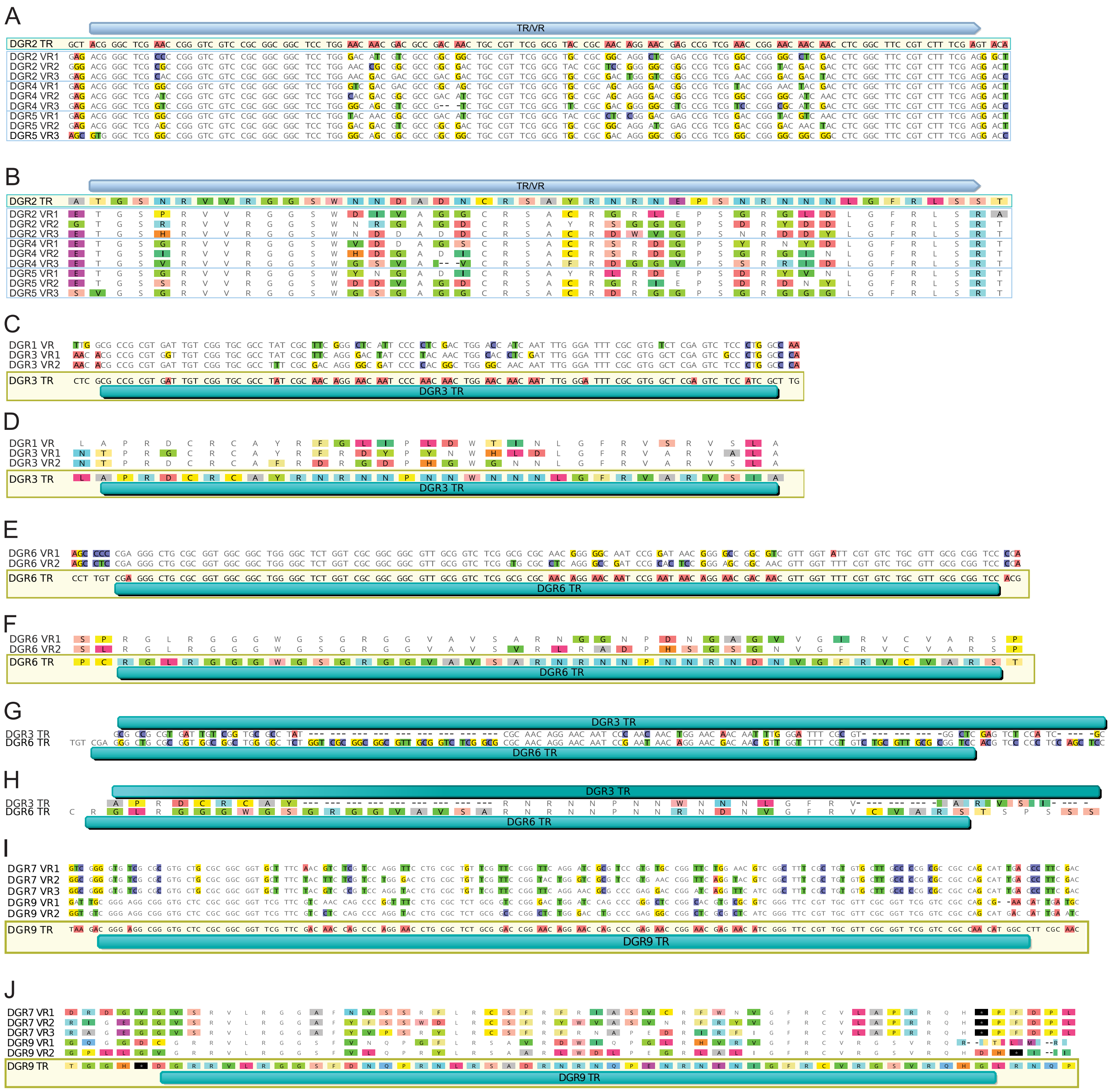

**Fig. S1.** Alignment of VRs-TR from DGR2-4-5, DGR1-3-6 and DGR7-9.
(**A**) Nucleotide alignment of the TR from DGR2 (top, highlighted in yellow), and the VRs from DGR2/4/5. The predicted TR/VR region is indicated above the sequences, and the positions that are As in the TR are shown in red. Only the mismatches to the TR are colored. (**B**) Amino-acid alignment of the TR from DGR2 and VRs from DGR2/4/5. Although the TR is not translated, the theoretical sequence is shown for comparison to the VRs. (**C**) Nucleotide alignment of the TR of DGR3 and the VRs of DGR1 and DGR3. The location of the predicted TR region is indicated below the sequences, and the positions that are As in the TR are shown in red. Only the mismatches to the TR are colored. (**D**) Amino acid alignment of the TR of DGR3 and the VRs from DGR1 and DGR3. Although the TR is not translated, the theoretical sequence is shown for comparison to the VRs. (**E**) Nucleotide alignment of the TR and VRs of DGR6. The location of the predicted TR region is indicated below the sequences, and the positions that are As in the TR are shown in red. Only the mismatches to the TR are colored. (**F**) Amino acid alignment of the TR and VRs from DGR6. Although the TR is not translated, the theoretical sequence is shown for comparison to the VRs. (**G**) Nucleotide alignment of DGR3 TR and DGR6 TR. Differences between the two sequences are highlighted. (**H**) Amino-acid alignment of DGR3 TR and DGR6 TR. Although the TR is not translated, the theoretical sequence is shown to show the effect of substitutions. (**I**) Nucleotide alignment of the TR from DGR9 (bottom, highlighted in yellow), and the VRs from DGR7/9. The predicted TR/VR region is indicated below the sequences, and the positions that are As in the TR are shown in red. Only the mismatches to the TR are colored. (**J**) Amino-acid alignment of the TR from DGR9 and VRs from DGR7/9. Although the TR is not translated, the theoretical sequence is shown for comparison to the VRs.

A

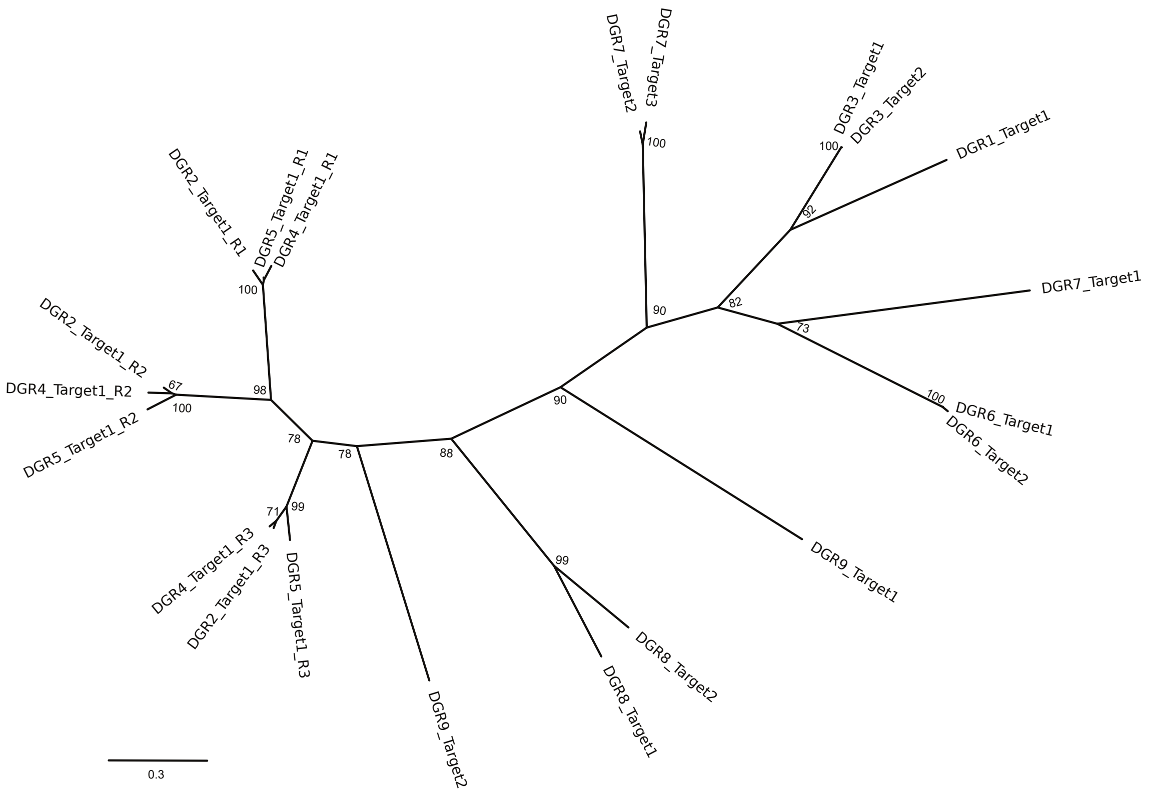


B

| | DGR2_Target1_R1 | DGR5_Target1_R1 | DGR4_Target1_R1 | DGR2_Target1_R2 | DGR4_Target1_R2 | DGR5_Target1_R2 | DGR2_Target1_R3 | DGR4_Target1_R3 | DGR5_Target1_R3 | DGR9_Target2 | DGR8_Target1 | DGR8_Target2 | DGR3_Target1 | DGR3_Target2 | DGR1_Target1 | DGR7_Target1 | DGR6_Target2 | DGR6_Target1 | DGR7_Target2 | DGR7_Target3 | DGR9_Target1 |
|---|---|---|---|---|---|---|---|---|---|---|---|---|---|---|---|---|---|---|---|---|---|
| DGR2_Target1_R1 | | 96 | 70 | 70 | 70 | 63 | 63 | 61 | 48 | 41 | 42 | 40 | 40 | 40 | 43 | 38 | 38 | 42 | 43 | 40 | 40 |
| DGR5_Target1_R1 | 97 | | 96 | 70 | 71 | 70 | 63 | 64 | 62 | 48 | 41 | 42 | 39 | 39 | 40 | 43 | 38 | 38 | 42 | 43 | 39 |
| DGR4_Target1_R1 | 96 | 96 | | 70 | 70 | 70 | 63 | 64 | 62 | 48 | 40 | 42 | 40 | 40 | 40 | 43 | 37 | 37 | 42 | 43 | 38 |
| DGR2_Target1_R2 | 70 | 70 | 70 | | 98 | 94 | 67 | 68 | 65 | 51 | 46 | 49 | 37 | 37 | 38 | 43 | 39 | 39 | 41 | 41 | 44 |
| DGR4_Target1_R2 | 70 | 71 | 70 | 98 | | 93 | 66 | 68 | 65 | 52 | 46 | 50 | 38 | 38 | 38 | 43 | 40 | 40 | 40 | 40 | 44 |
| DGR5_Target1_R2 | 70 | 70 | 70 | 94 | 93 | | 65 | 67 | 60 | 50 | 44 | 48 | 37 | 37 | 38 | 41 | 40 | 40 | 40 | 40 | 42 |
| DGR2_Target1_R3 | 63 | 63 | 63 | 67 | 66 | 65 | | 95 | 85 | 46 | 46 | 45 | 41 | 41 | 39 | 40 | 39 | 39 | 41 | 43 | 38 |
| DGR4_Target1_R3 | 63 | 64 | 64 | 68 | 68 | 67 | 95 | | 86 | 46 | 45 | 43 | 40 | 40 | 37 | 39 | 37 | 36 | 40 | 41 | 37 |
| DGR5_Target1_R3 | 61 | 62 | 62 | 65 | 65 | 60 | 85 | 86 | | 46 | 46 | 42 | 38 | 38 | 35 | 39 | 35 | 35 | 40 | 41 | 37 |
| DGR9_Target2 | 48 | 48 | 48 | 51 | 52 | 50 | 46 | 46 | 46 | | 51 | 53 | 35 | 35 | 33 | 39 | 40 | 40 | 48 | 47 | 43 |
| DGR8_Target1 | 41 | 41 | 40 | 46 | 46 | 44 | 46 | 45 | 46 | 51 | | 77 | 33 | 33 | 31 | 38 | 41 | 41 | 41 | 42 | 45 |
| DGR8_Target2 | 42 | 42 | 42 | 49 | 50 | 48 | 45 | 43 | 42 | 53 | 77 | | 34 | 34 | 34 | 40 | 43 | 43 | 40 | 42 | 45 |
| DGR3_Target1 | 40 | 39 | 40 | 37 | 38 | 37 | 41 | 40 | 38 | 35 | 33 | 34 | | 100 | 59 | 41 | 42 | 42 | 49 | 49 | 33 |
| DGR3_Target2 | 40 | 39 | 40 | 37 | 38 | 37 | 41 | 40 | 38 | 35 | 33 | 34 | 100 | | 59 | 41 | 42 | 42 | 49 | 49 | 33 |
| DGR1_Target1 | 40 | 40 | 40 | 38 | 38 | 38 | 39 | 37 | 35 | 33 | 31 | 34 | 59 | 59 | | 43 | 47 | 47 | 48 | 48 | 32 |
| DGR7_Target1 | 43 | 43 | 43 | 43 | 43 | 41 | 40 | 39 | 39 | 39 | 38 | 40 | 41 | 41 | 43 | | 54 | 54 | 45 | 44 | 37 |
| DGR6_Target2 | 38 | 38 | 37 | 39 | 40 | 40 | 39 | 37 | 35 | 40 | 41 | 43 | 42 | 42 | 47 | 54 | | 99 | 54 | 54 | 41 |
| DGR6_Target1 | 38 | 38 | 37 | 39 | 40 | 40 | 39 | 36 | 35 | 40 | 41 | 43 | 42 | 42 | 47 | 54 | 99 | | 54 | 54 | 40 |
| DGR7_Target2 | 42 | 42 | 42 | 41 | 40 | 40 | 41 | 40 | 40 | 48 | 41 | 40 | 49 | 49 | 48 | 45 | 54 | 54 | | 96 | 42 |
| DGR7_Target3 | 43 | 43 | 43 | 41 | 40 | 40 | 43 | 41 | 41 | 47 | 42 | 42 | 49 | 49 | 48 | 44 | 54 | 54 | 96 | | 42 |
| DGR9_Target1 | 40 | 39 | 38 | 44 | 44 | 42 | 38 | 37 | 37 | 43 | 45 | 45 | 33 | 33 | 32 | 37 | 41 | 40 | 42 | 42 | |

**Fig. S2.** Sequence similarity of the C-terminal lectin domains from *Thiohalocapsa* sp. PB-PSB1's 15 DGR target proteins.

(**A**) Maximum likelihood phylogeny of the CLec domains from the 15 target proteins. The variable region of each CLec domain has been removed prior to alignment. Bootstrap support is shown at each node (n=100). Each domain is identified by the DGR locus, target gene name, and for targets with multiple CLec domains they have been numbered from 5' to 3' (R1, R2, R3). (**B**) Amino acid similarity (BLOSUM 62) distance matrix for the CLec domain alignment used for the phylogeny in panel A (the variable region of each CLec domain has been removed prior to alignment).

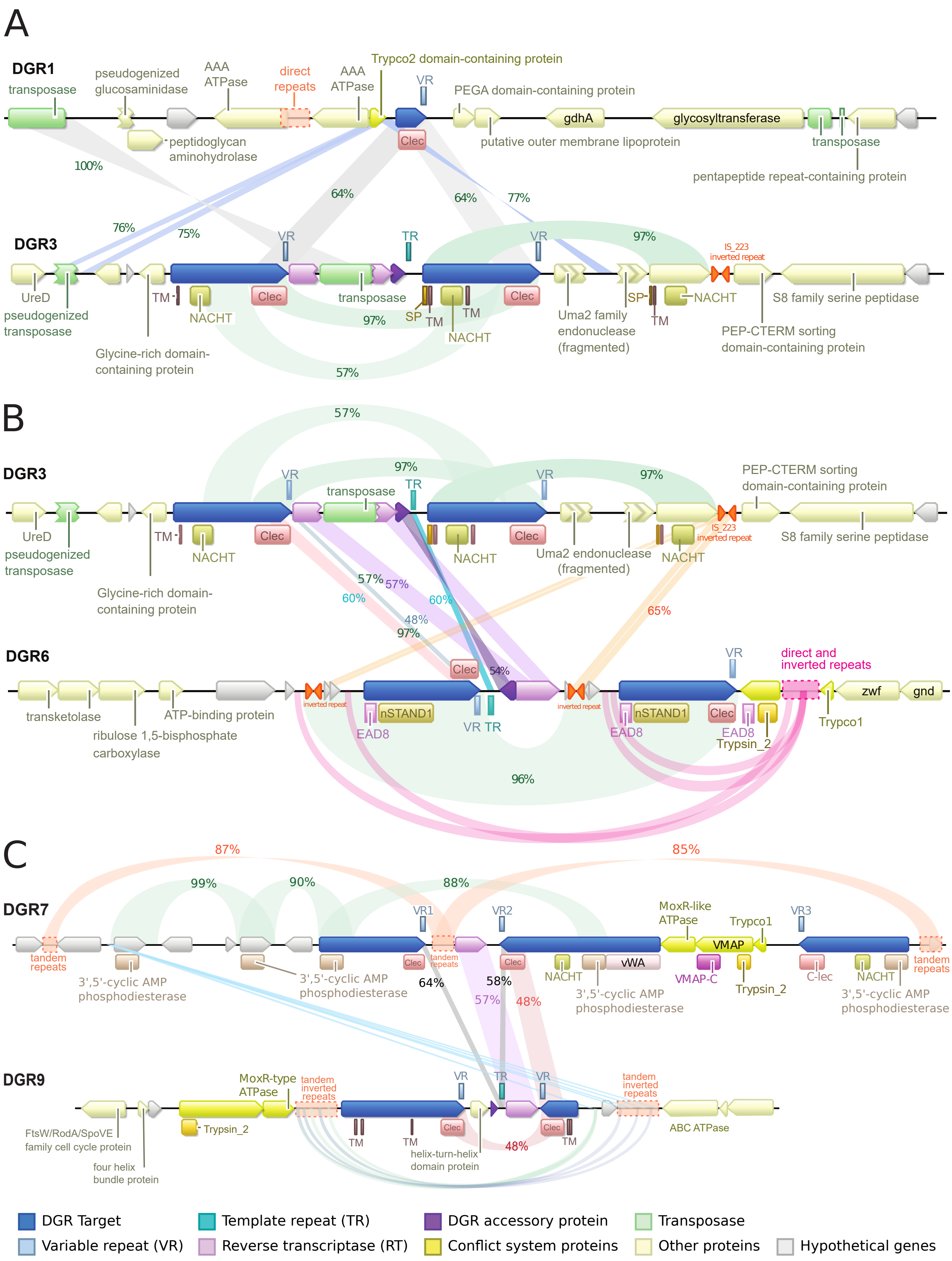


**Fig. S3.** Comparison of DGR loci 1 and 3 (**A**), 3 and 6 (**B**) and 7 and 9 (**C**). The gene neighborhoods surrounding each DGR locus are shown with regions of similarity highlighted along with the percent nucleotide identity.

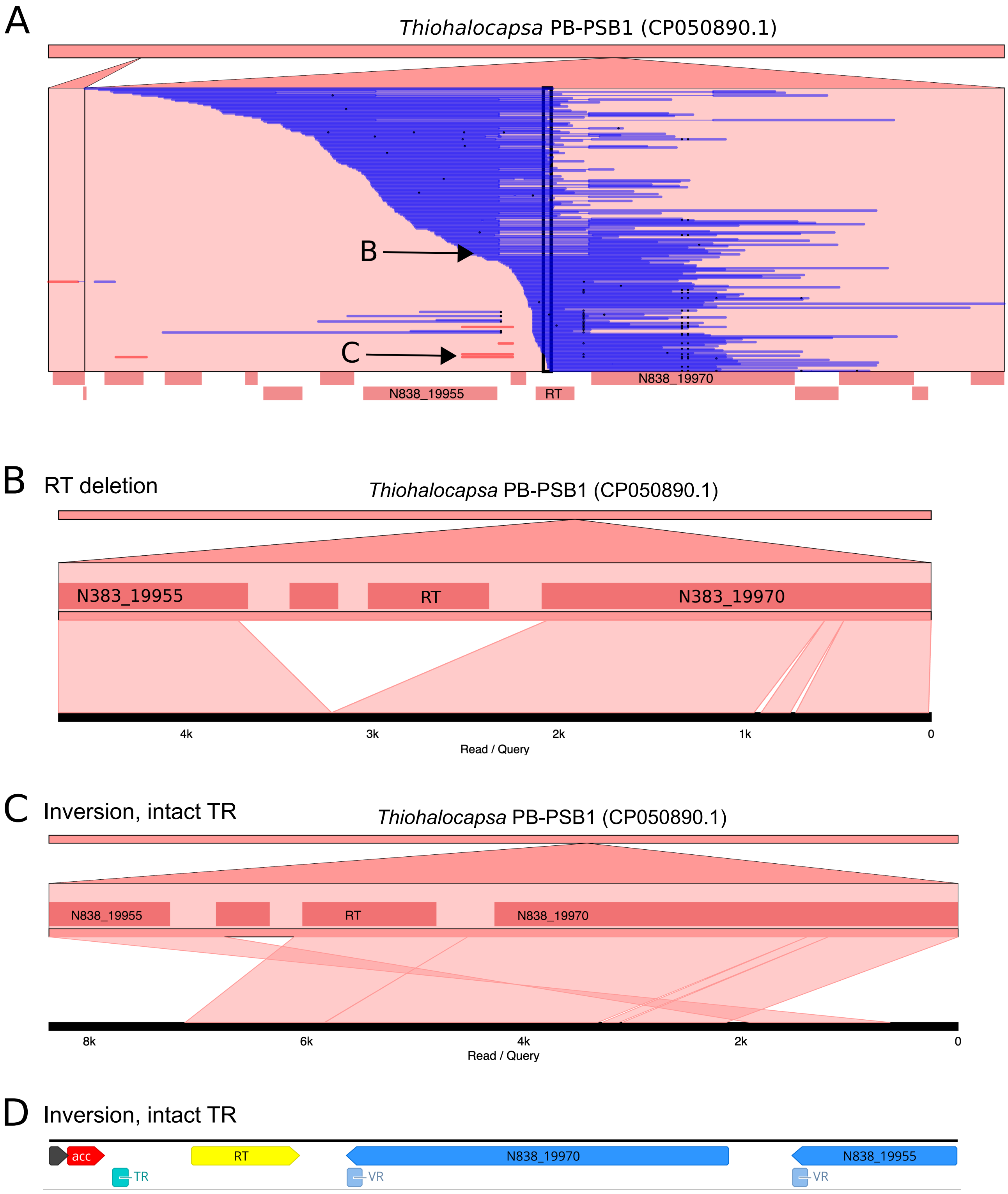


**Fig. S4.** Structural variants observed at the DGR7 locus.
Structural variants are shown as seen in the long-read alignments to the *Thiohalocapsa* sp. PB-PSB1 reference genome visualized with Genome Ribbon (5). (**A**) Read mappings are shown in blue, with inversions shown in red, short indels shown in black, and longer deletions shown with a thin blue connecting line. Ribbon plots of individual reads show examples of variants with a deleted RT gene (**B**) and variants with inversions and an intact template repeat region (TR) (**C**). Panel (**D**) shows the same read as in panel (**C**) with the intact DGR region annotated using myDGR (6).

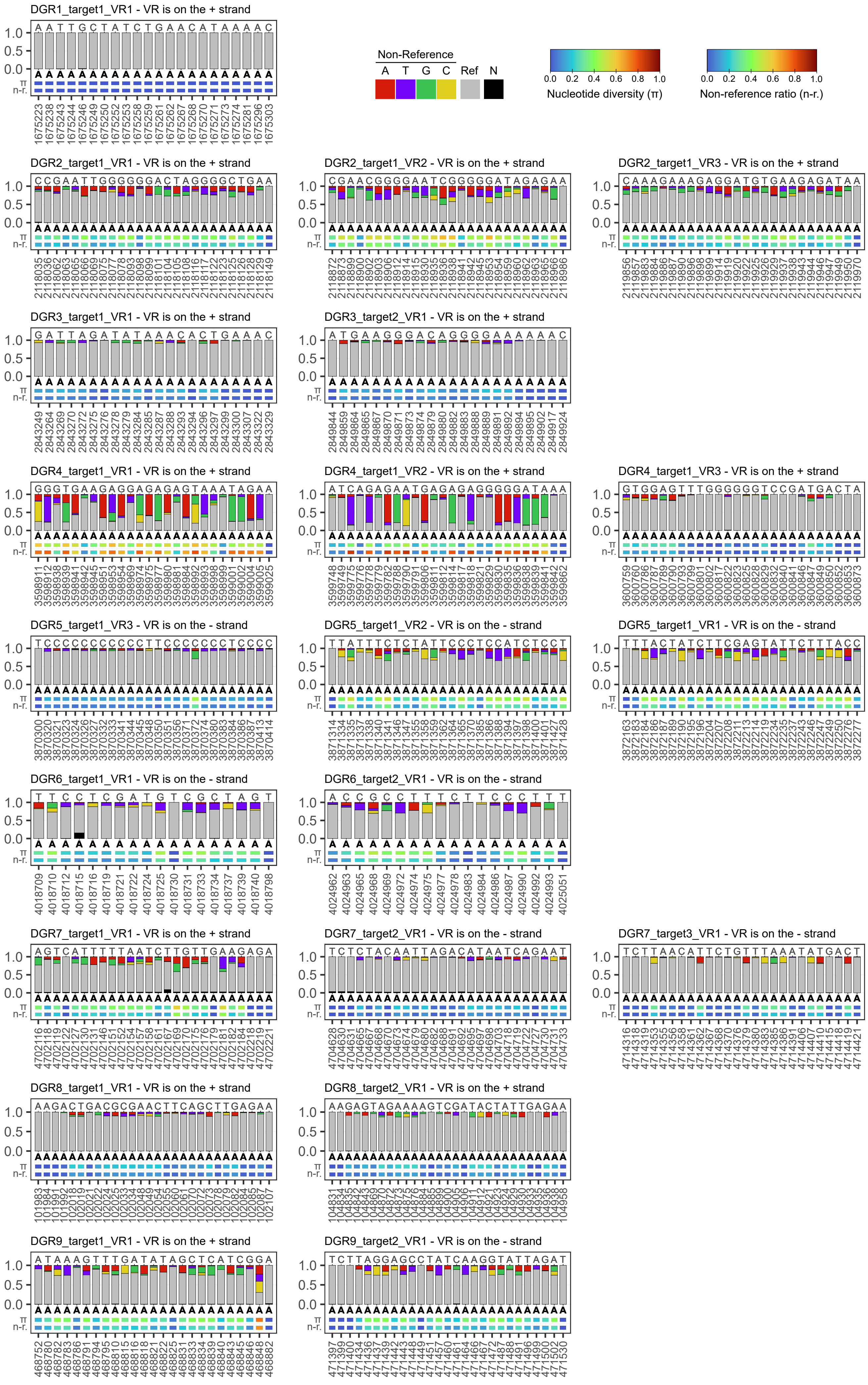

**Fig. S5.** *In situ* diversification of VRs of all DGR loci in LS01_001 pink berry aggregate from long-reads metagenomics data.
Only positions that correspond to an A in the TR are shown. Bar plots indicate the proportion of A, T, C and G at each position, colored if they differ from the reference. Letters above bars indicate the VR sequence in the reference genome, while letters below bars indicate the reference sequence of the TR. Bottom rows show the nucleotide diversity (π) and proportion of non-reference alleles (n-r.) at each position. Ref.: Reference nucleotide. N: unknown nucleotide.

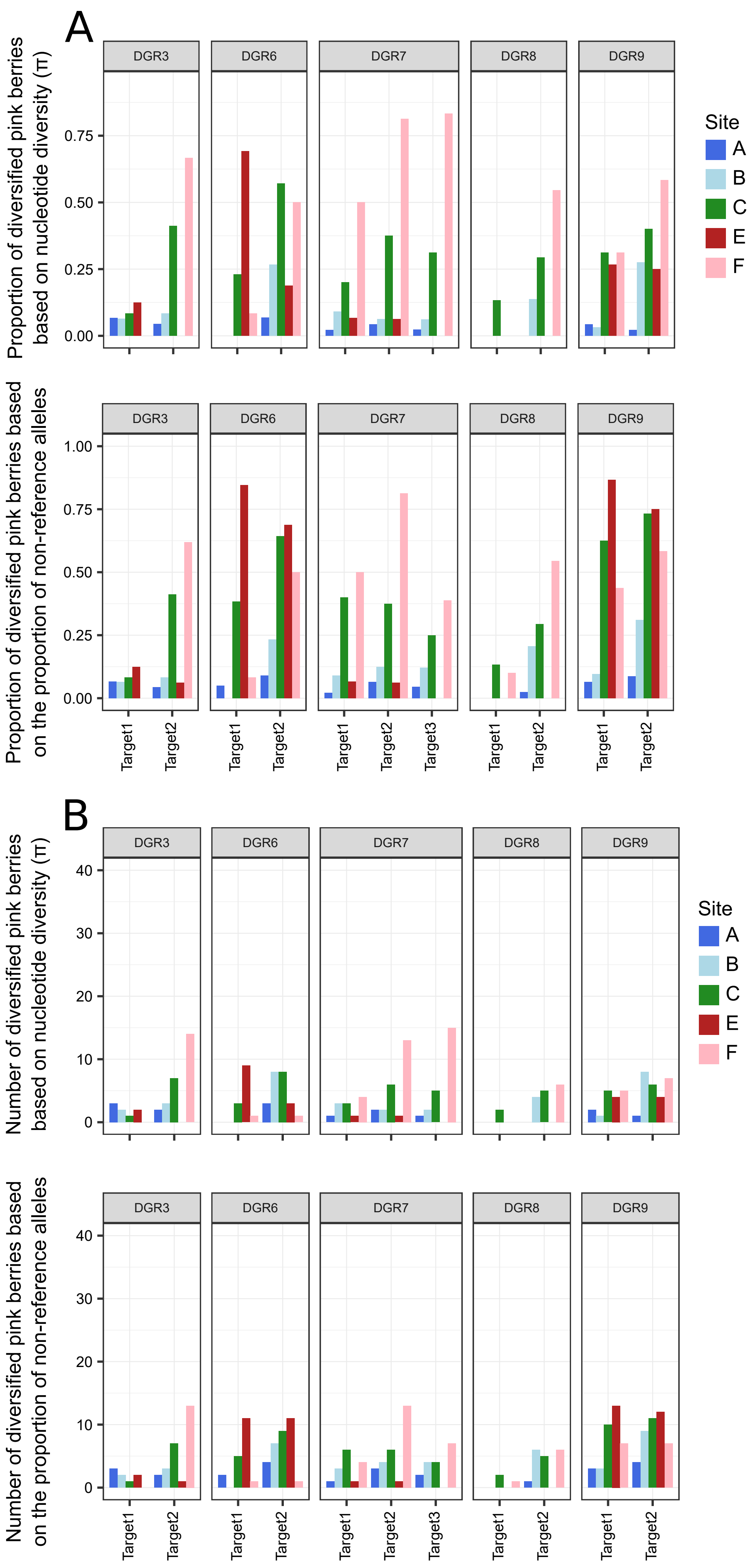


**Fig. S6.** Proportion (**A**) and number (**B**) of aggregates showing diversification for each DGR target gene at each sampling site. A DGR target was considered diversified if at least one of its VRs showed diversification at positions targeted by the DGR mechanism based on the nucleotide diversity (upper panel) or the proportion of non-reference alleles (lower panel). Only DGR loci with multiple targets are represented. Colors correspond to sampling sites, with blue shades corresponding to Little Sippewissett salt marsh (LS), green to Great Sippewissett (GS) and red shades to Penzance Point (PP).

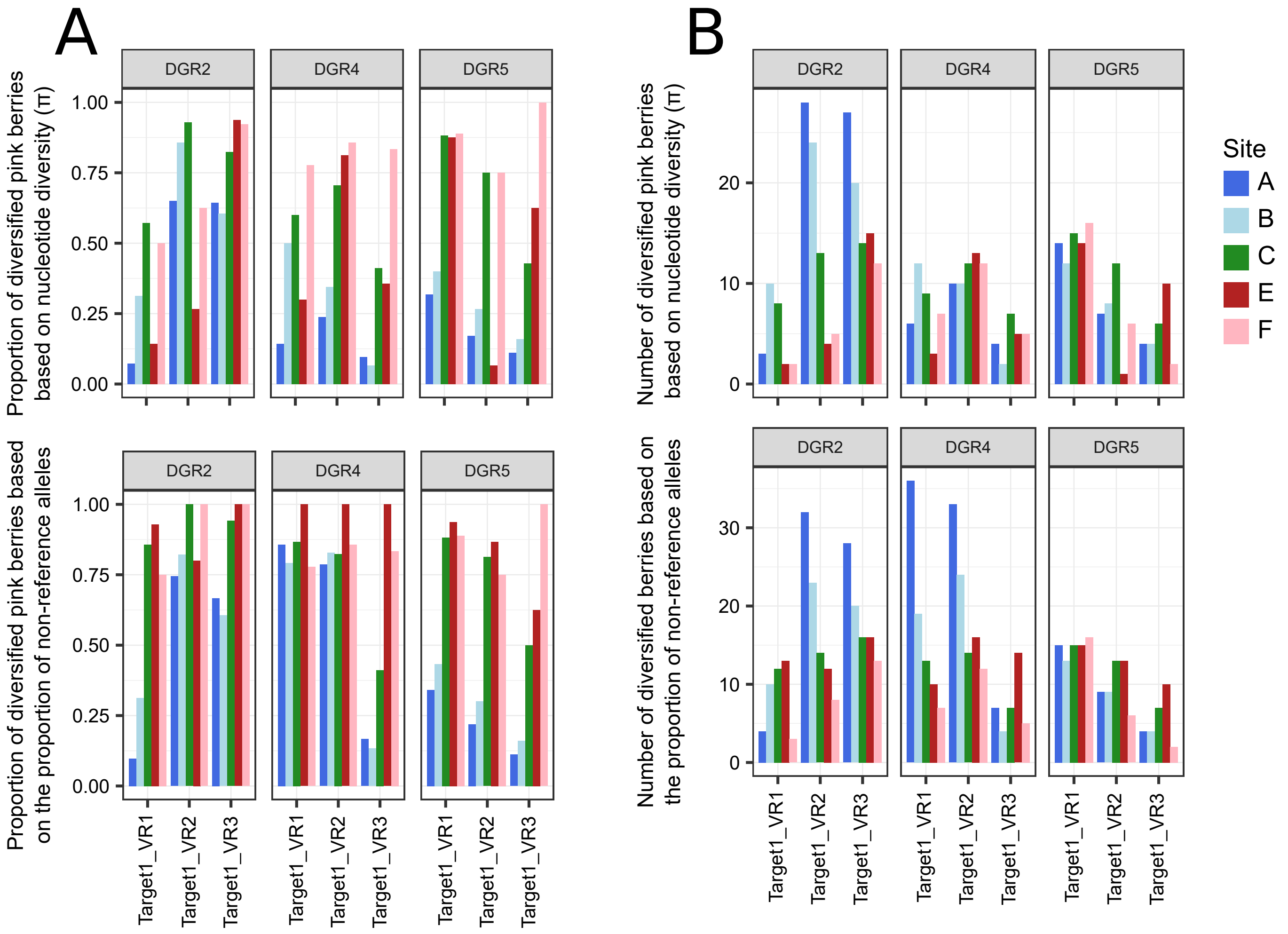


**Fig. S7.** Proportion (**A**) and number (**B**) of aggregates showing diversification for each VR of each DGR target gene at each sampling site. Nucleotide diversity (upper panel) or the proportion of non-reference alleles (lower panel) were used to determine if a VR was diversified. Only DGR targets with multiple VRs are represented. Colors correspond to sampling sites, with blue shades corresponding to Little Sippewissett salt marsh (LS), green to Great Sippewissett salt marsh (GS) and red shades to Penzance Point salt marsh (PP).

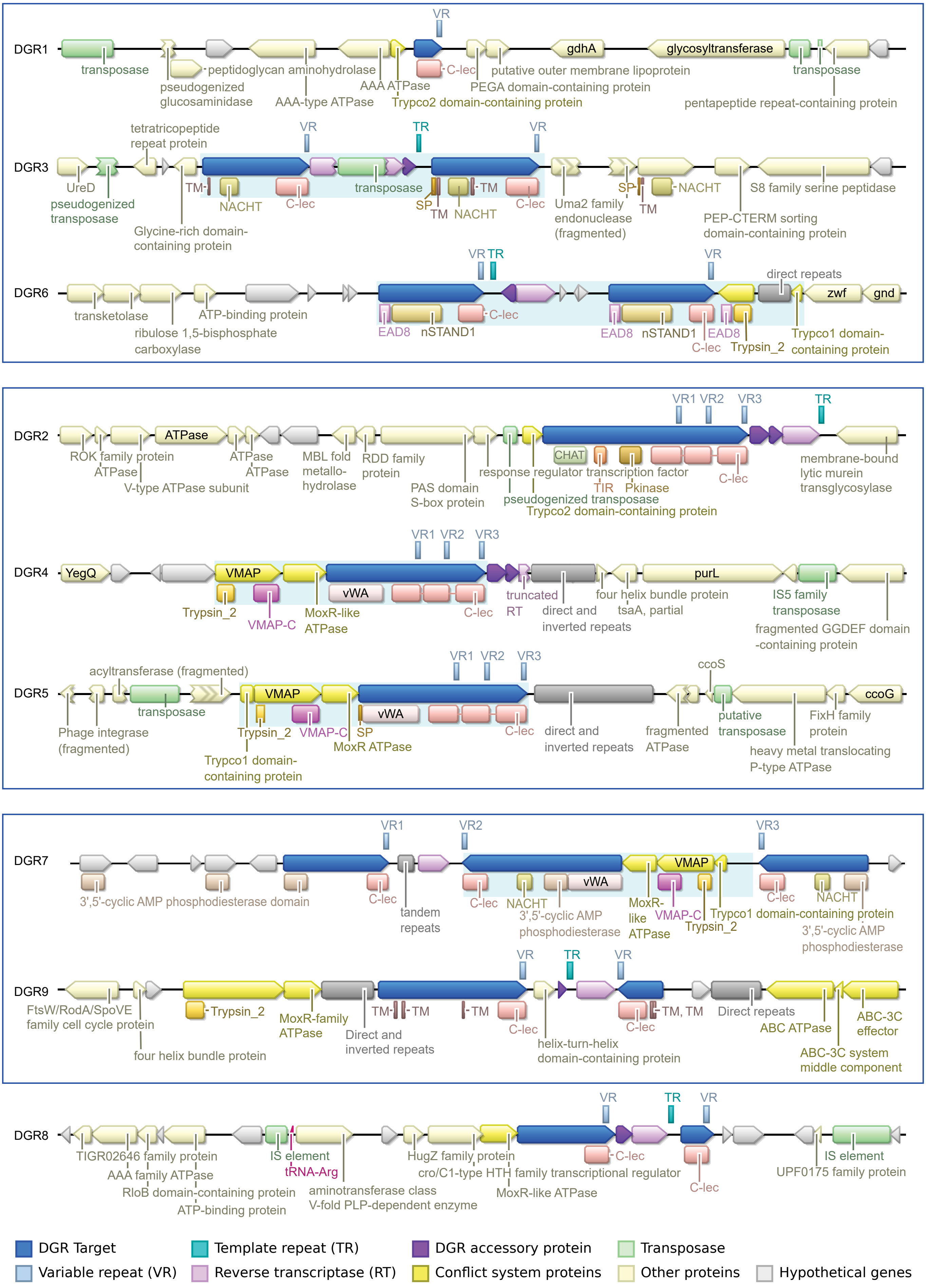


**Fig. S8.** Functional annotation of genes surrounding all *Thiohalocapsa* PB-PSB1 DGR loci.
The light blue shaded areas indicate gene architectures described as putative conflict systems. SP: signal peptide, TM: transmembrane domain.

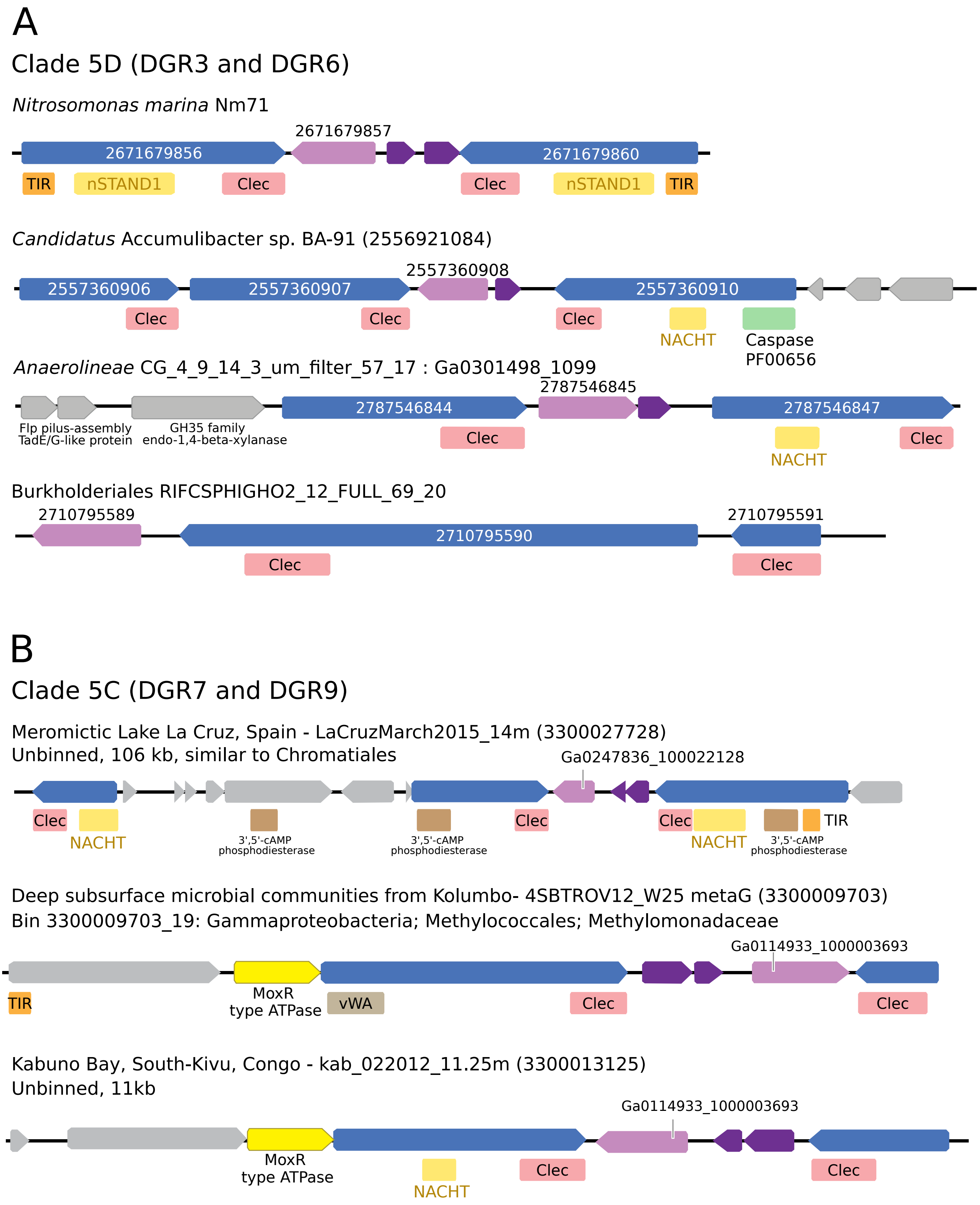


**Fig. S9.** Examples of clade 5 DGR loci with a gene organization similar to *Thiohalocapsa* PB-PSB1 DGRs. (**A**) DGR loci in clade 5D. (**B**) DGR loci in clade 5C. These examples correspond to metagenomic contigs. (**C**) DGR loci in clade 5B. The last example corresponds to a metagenomics contig. (**D**) DGR loci in clade 5A.

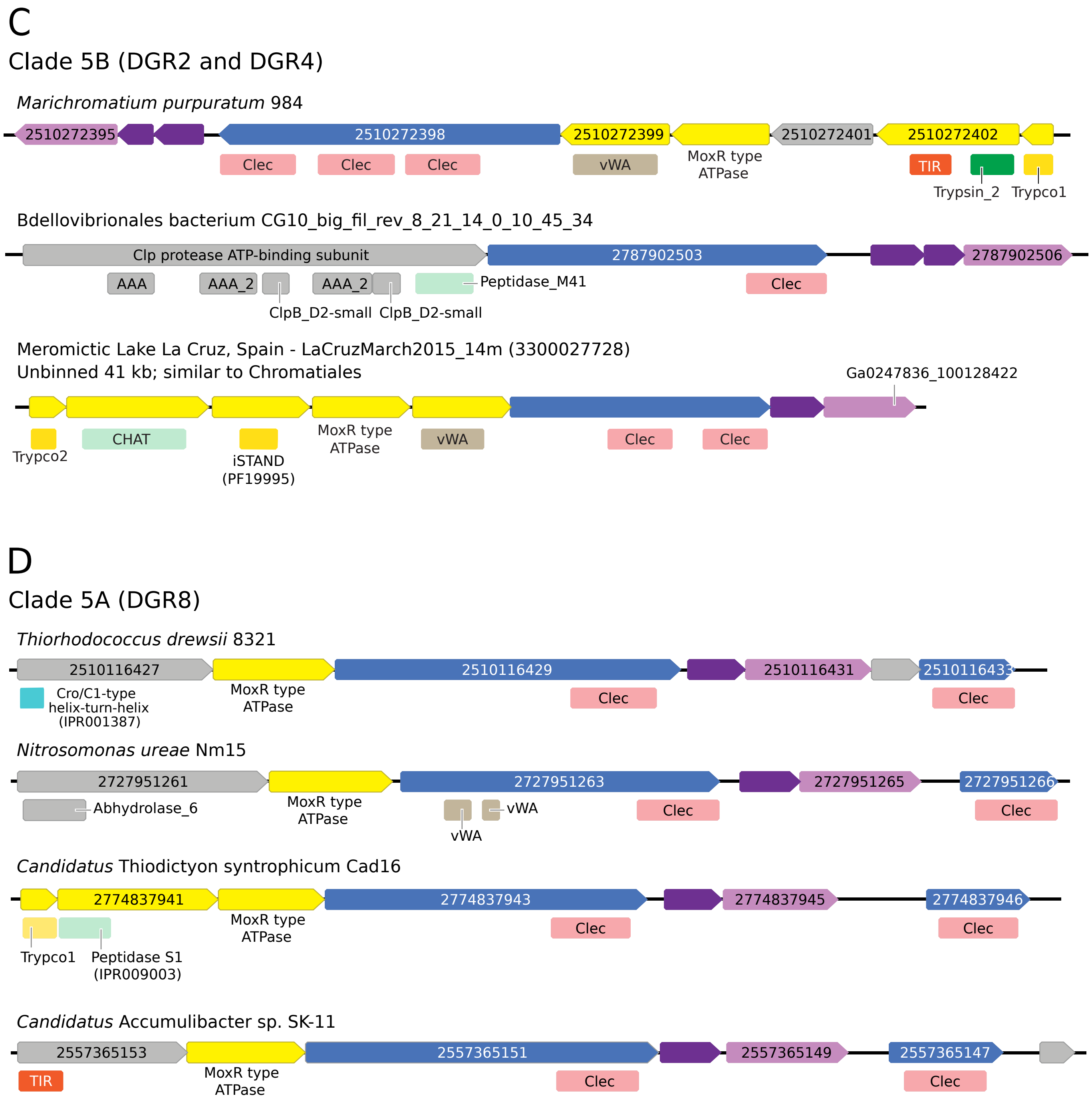


**Fig. S9.** Examples of clade 5 DGR loci with a gene organization similar to *Thiohalocapsa* PB-PSB1 DGRs. (**A**) DGR loci in clade 5D. (**B**) DGR loci in clade 5C. These examples correspond to metagenomic contigs. (**C**) DGR loci in clade 5B. The last example corresponds to a metagenomics contig. (**D**) DGR loci in clade 5A.
